# Prematurity insults remodel cerebellar development and behavior

**DOI:** 10.1101/2025.07.22.664624

**Authors:** Georgios Sanidas, Gabriele Simonti, Javid Ghaemmaghami, Kyla Woyshner, Robinson Vidva, Daniel Bittel, Nora Wolff, Maria Triantafyllou, Antonis Polyviou, Chad Byrd, Courtney Lowe, Henry Salisbury, Evan Goldstein, Aaron Sathyanesan, Ioannis Koutroulis, Genevieve Stein-O’Brien, Dimitrios N. Sidiropoulos, Vittorio Gallo, Panagiotis Kratimenos

**Affiliations:** Children’s National Research Institute, Children’s National Hospital, Washington, DC 20010, USA; Department of Pediatrics, The George Washington University School of Medicine and Health Sciences, Washington, DC 20037, USA; Department of Computational Medicine and Bioinformatics, University of Michigan, Ann Arbor, MI 48109, USA; Johns Hopkins University School of Medicine, Baltimore, MD 21205, USA; University of Dayton, Dayton, OH 45469, USA; Departments of Biological Sciences and Neuroscience & Regenerative Medicine, Augusta University, Augusta, GA 30912, USA; Seattle Children’s Research Institute, Seattle, WA 98101, USA; Department of Pediatrics, University of Washington School of Medicine, Seattle, WA 98195, USA

**Keywords:** Maternal immune activation, hypoxia, cerebellum, prematurity, locomotor, social behavior

## Abstract

Preterm survivors often develop motor and socio-cognitive impairments that implicate altered cerebellar development, yet the underlying mechanisms remain poorly understood. A key challenge is that prematurity involves overlapping perinatal insults that converge on the developing brain, making their individual effects difficult to disentangle.

Here, we model two major prematurity-associated insults, maternal immune activation (MIA) and neonatal hypoxia (Hx), in mice. By controlling timing and sequence, we define how these insults shape cerebellar assembly during rapid maturation. Comparative analysis with human tissue confirmed that these insults recapitulate key features of the human preterm cerebellum, establishing the translational validity of this model for dissecting insult-specific outcomes.

Behavioral and kinematic profiling revealed divergent motor and social phenotypes, which we traced to insult-specific cerebellar remodelling. Hypoxia compromised both granule cell maturation in the internal granule layer and the principal excitatory afferents to Purkinje cells, a circuit state that manifested as impaired motor execution and stereotyped, sensory-disengaged social investigation. Maternal immune activation, by contrast, expanded the granule cell progenitor layer and reduced Purkinje cell dendritic complexity, a phenotype that preserved social preference but reorganized the kinematic structure of social investigation. When hypoxia followed maternal immune activation, it acted on this primed substrate to produce a distinct state in which the features established by prior inflammation were compounded by granule cell proliferative arrest, progressive mitochondrial dysfunction, and aberrant, hyper-interactive social investigation.

Together, these findings reframe prematurity-associated insults as cerebellar reprogramming events shaped by both the identity and sequence of insults, linking distinct mechanistic substrates to divergent neurodevelopmental outcomes.

## Introduction

Despite major advances in neonatal intensive care that have improved survival among extremely preterm infants (<28 weeks’ gestation)^1,2,3^, the prevalence of long-term neurodevelopmental impairment has not declined^3^. Preterm birth exposes the developing brain to a constellation of biologically active insults that disrupt key neurodevelopmental processes^4,5^.

Among the most prevalent prenatal contributors is maternal immune activation (MIA), which is implicated in a large fraction of preterm births associated with intrauterine infection or inflammation^6^. MIA perturbs the tightly regulated immune environment required to maintain pregnancy and disrupts fetal neurodevelopmental programs^7,8,9^, increasing vulnerability to later socioemotional and cognitive impairments^10,11^.

Postnatally, hypoxic exposure is one of the most frequent and consequential insults in prematurity^12^. Lung immaturity often necessitates prolonged ventilatory support^13^, and early-life hypoxia disrupts neuronal differentiation, synaptogenesis, and myelination, resulting in long-term behavioral deficits^14,15^.

The cerebellum is particularly vulnerable in this context. Its abrupt transition to the *ex-utero* environment during a critical window of maturation increases sensitivity to inflammatory and hypoxic stressors that can alter structural integrity and circuit formation^16^. Disruption of these developmental processes has functional consequences that extend beyond motor control, with cerebellar dysfunction implicated in ataxia as well as socioemotional and cognitive deficits^2^ and increasingly linked to neurodevelopmental conditions, such as autism spectrum disorder^17,18^. Yet the mechanistic interplay between prematurity-related insults and cerebellar development remains incompletely understood.

Here, we resolve how MIA and Hx, both individually and sequentially, shape cerebellar circuit assembly, and how they differentially alter motor learning, coordination, and social behavior. We utilized a temporally controlled neonatal mouse model to establish how isolated and sequential prematurity-associated insults differentially drive cerebellar-mediated behavioral abnormalities. We then applied spatial transcriptomic and histological analyses to trace these functional deficits back to their structural and bioenergetic roots, resolving the divergent cellular mechanisms underlying each phenotype. We demonstrate how MIA and hypoxia, isolated and in sequence, disrupt distinct elements of cerebellar maturation, including granule cell development, Purkinje cell integrity, and cerebellar input pathways, a configuration whose molecular state most closely recapitulates the human preterm cerebellum. Together, these findings reframe prematurity-associated cerebellar injury not as a uniform pathology, but as a dynamic process where the exact identity and sequence of early-life insults dictate circuit assembly and the resulting behavioral phenotypes.

## Results

### Experimental design of prematurity-associated insults

To test how prematurity-related insults shape long-term neurobehavioral outcomes, we used an experimental neonatal mouse model that allows controlled comparison of isolated and sequential exposures. Because spontaneous preterm labor in humans is frequently triggered by maternal bacterial infection^49,50^, prenatal MIA was induced using lipopolysaccharide (LPS), a Toll-like receptor 4 agonist that recapitulates bacterially driven inflammatory signaling^51,20^. LPS was administered at embryonic days 15–16 (E15–16), a developmental window corresponding to 22–24 weeks of human gestation, to induce inflammation-mediated spontaneous preterm birth, which consistently occurred at E18–19^49,52,6,53^(detailed in *Materials and Methods*).

We modeled respiratory insufficiency to mimic preterm pulmonary immaturity using a validated neonatal sublethal chronic hypoxia paradigm (10.5-11% FiO_2_)^54,55^. Pups were exposed to hypoxia across the first postnatal week (P3 to P11), approximating the cumulative hypoxic burden experienced by extremely preterm infants in the NICU^56^ (see details in *Materials and Methods*). Long-term functional outcomes were evaluated through motor and socio-behavioral testing in adulthood (Fig. 1A). We included four groups across insult conditions: saline-normoxia controls (Sal-N), isolated MIA (LPS-N), isolated hypoxia (Sal-Hx), and sequential exposure (LPS-Hx).

**Fig. 1.**
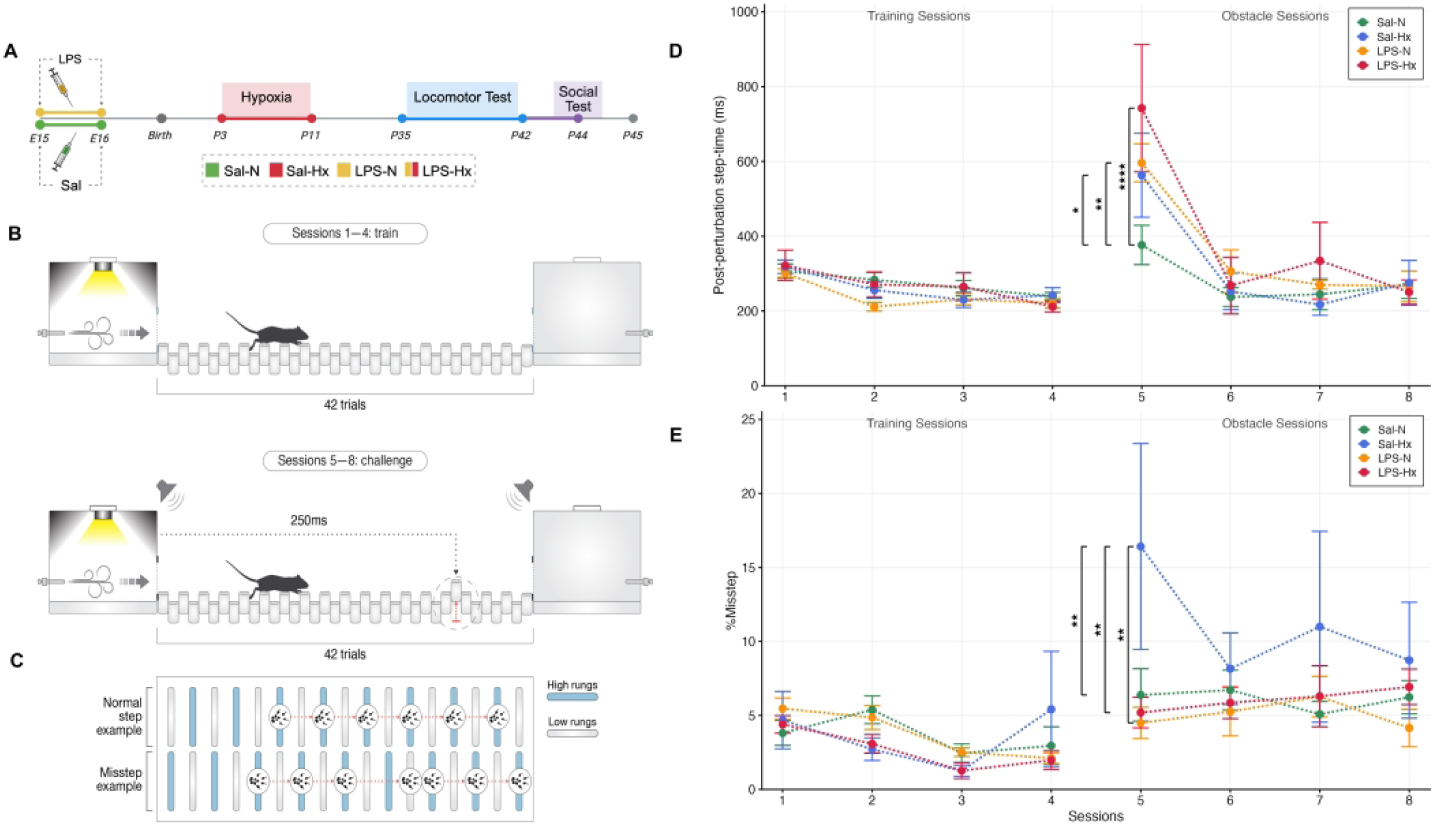
Distinct Long-Term Cerebellar Motor Deficits Induced by MIA and Hx. **(A)** Experimental timeline of the sequential prematurity insult mouse model. Pregnant dams received intraperitoneal injections of lipopolysaccharide (LPS) or saline (Sal) at embryonic days 15 and 16 (E15-E16) to induce maternal immune activation. Offspring were subsequently exposed to chronic hypoxia (Hx) or maintained under normoxic conditions (N) from postnatal day 3 through 11 (P3-P11). Cerebellar-dependent motor function was evaluated using the ErasmusLadder at P35-P42, followed by social behavior testing at P44 ± 2 days at the time of testing. **(B)** Schematic of the ErasmusLadder paradigm. Mice traversed a horizontal ladder from an origin box to a goal box (equivalent of one trial) over eight sessions of 42 trials each. Sessions 1-4 served as training under unperturbed conditions, whereas sessions 5-8 introduced challenge trials incorporating an auditory cue, an obstacle (raised rung), or both, presented in randomized order. **(C)** Representative diagrams illustrate normal stepping patterns and missteps, defined as limb placement errors reflecting impaired motor coordination. **(D)** Post-perturbation step-time across sessions. All groups performed comparably during training sessions. Upon obstacle introduction, Sal-Hx, LPS-N, and LPS-Hx mice exhibited significantly prolonged step-times relative to Sal-N controls (Sal-N versus Sal-Hx, \**P* = 0.025; Sal-N versus LPS-N, \*\**P* = 0.0054; Sal-N versus LPS-Hx, \*\*\*\**P* < 0.0001). **(E)** Misstep frequency across sessions. Sal-Hx mice demonstrated elevated misstep rates during obstacle sessions compared with all other groups (Sal-Hx versus Sal-N, \*\**P* = 0.0023; Sal-Hx versus LPS-N, \*\**P* = 0.0011; Sal-Hx versus LPS-Hx, \*\**P* = 0.0023). Data are presented as mean ± SEM; *n* = 6-10 animals per group; two-way repeated-measures ANOVA with Tukey’s post hoc correction for multiple comparisons; \**P* < 0.05, \*\**P* < 0.01, \*\*\**P* < 0.001, \*\*\*\**P* < 0.0001. Abbreviations: E, embryonic day; Hx, hypoxia; LPS, lipopolysaccharide; N, normoxia; P, postnatal day; Sal, saline; SEM, standard error of the mean.

### Prematurity-associated insults differentially disrupt cerebellar motor-learning and coordination

Preterm-born children often exhibit ataxia and impaired coordination, pointing to persistent cerebellar dysfunction^57^. To determine how prematurity-related insults shape cerebellar-dependent behavior, we used the Erasmus Ladder, a high-resolution assay of locomotor learning, adaptive timing, and gait precision, across four training sessions followed by four obstacle-challenge sessions (42 trials per session, Fig. 1B-C)^58,37^.

As repeated exposure to the obstacle (Sessions 5 to 8) progressively elicits motor adaptation, the magnitude of the latency to approach the obstacle at challenge onset indicates the degree of impaired sensorimotor adaptation (Fig. 1D, Supplementary Table 9A). In the initial challenge phase (Session 5), each insult group showed delayed obstacle responses relative to Sal-N controls. This deficit was most severe in the sequential LPS-Hx group, which exhibited a nearly two-fold increase in response latency (Fig. 1D).

In the domain of pure motor coordination, measured by misstep frequency (Fig. 1C, E), isolated hypoxia exerted a pronounced and selective deficit in motor execution and gait stability with a threefold increase in missteps compared to all other groups, including those exposed to both insults (Fig. 1E, Supplementary Table 9B). Notably, this coordination deficit was entirely absent in both the isolated MIA (LPS-N) group and the sequential insult (LPS-Hx) group, both of which maintained normal baseline misstep rates.

Together, these results indicate that perinatal insults drive separable motor trajectories rather than a cumulative injury response. Hypoxia alone preferentially impaired motor execution and gait stability, whereas sequential LPS-Hx exposure produced the strongest disruption of adaptive motor learning. Thus, motor outcomes were determined by exposure identity and sequence, indicating that distinct perinatal insults selectively compromise different functional cerebellar circuits.

### Sequential MIA and Hx recapitulate features of the preterm human cerebellum

To resolve how prematurity-related insults shape the cerebellum across development; we examined cerebellar pathology at early postnatal and adult stages (Fig. 2A). In MIA-exposed mice, irrespective of subsequent Hx, Purkinje cell density was significantly reduced in early life at P11 compared to controls (Fig. 2B). However, these changes were no longer detectable by adulthood at P45 (Fig. 2C). By contrast, granule cell (GC) lineage disruption persisted into adulthood. At P11, MIA exposure resulted in expansion of the external granular layer (EGL) in both the LPS-N and LPS-Hx groups relative to Sal-N controls (Fig. 2D), indicating perturbed granule cell precursor dynamics through altered progenitor cell-cycle and migration states. Additionally, and irrespective of MIA, hypoxia selectively reduced GC density in the internal granule layer (IGL) at both P11 and P45 (Fig. 2E-F).

**Fig. 2.**
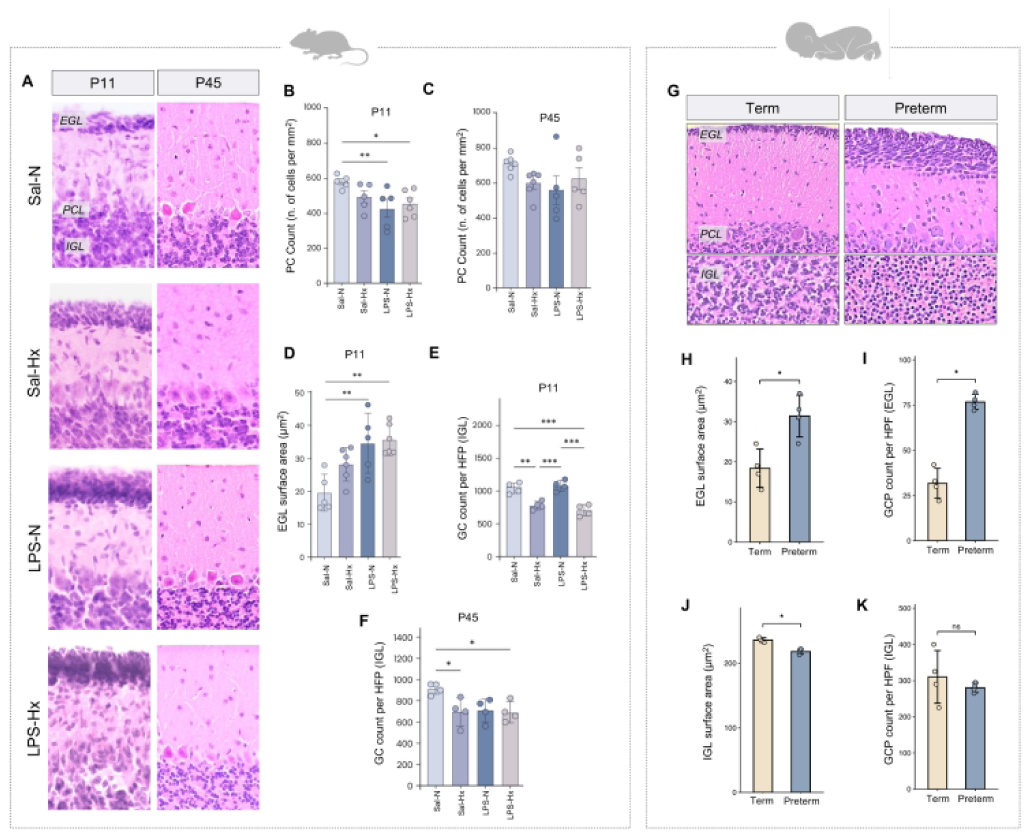
Prematurity Insults Disrupt Cerebellar Cytoarchitecture and Mimic Human Preterm Pathology. **(A)** Representative hematoxylin and eosin-stained photomicrographs of mouse cerebellar tissue at P11 and P45 across all four experimental groups (Sal-N, Sal-Hx, LPS-N, LPS-Hx), acquired at 60× magnification. **(B–C)** Purkinje cell count (number of cells per mm²) quantified at P11 (B) (Sal-N versus LPS-N, \*\**P* = 0.0082; Sal-N versus LPS-Hx, \**P* = 0.020) and P45 (C). **(D)** External granule layer surface area (µm²) measured at P11 (Sal-N versus LPS-N, \*\**P* = 0.0060; Sal-N versus LPS-Hx, \*\**P* = 0.0023). **(E)** Granule cell count per high-power field within the internal granule layer at P11 (Sal-N versus Sal-Hx, **P = 0.0025; Sal-N versus LPS-Hx, \*\*\**P* = 3.0 10⁻⁴; Sal-Hx versus LPS-N, \*\*\**P* = 8.0 × 10⁻⁴; LPS-N versus LPS-Hx, \*\*\**P* = 1.0 × 10⁻⁴). **(F)** Granule cell count per high-power field within the internal granule layer at P45 (Sal-N versus Sal-Hx, \**P* = 0.0497; Sal-N versus LPS-Hx, \**P* = 0.048). **(G–K)** Histological analysis of human cerebellar tissue from term and preterm infants. Representative photomicrographs (G) display the external granule layer, Purkinje cell layer, and internal granule layer in term and preterm cerebellum at 60× magnification. External granule layer surface area (H) (\**P* = 0.021) and granule cell precursor count per high-power field in the external granule layer (I) (\**P* = 0.021) in term versus preterm cerebellum. Internal granule layer surface area (J) (\**P* = 0.021) and granule cell precursor count per high-power field in the internal granule layer (K) (ns) in term versus preterm cerebellum. For mouse studies, *n* = 4–6 per group; one-way ANOVA with uncorrected Fisher’s LSD (B–C) or Tukey’s post hoc correction (D–F). For human studies (H, K) *n* = 4 term and *n* = 4 preterm subjects; two-tailed Mann–Whitney U test. Data are presented as mean ± SEM; \**P* < 0.05, \*\**P* < 0.01, \*\*\**P* < 0.001. Abbreviations: EGL, external granule layer; GC, granule cell; GCP, granule cell precursor; H&E, hematoxylin and eosin; HPF, high-power field; Hx, hypoxia; IGL, internal granule layer; LPS, lipopolysaccharide; N, normoxia; ns, not significant; P, postnatal day; PC, Purkinje cell; PCL, Purkinje cell layer; Sal, saline; SEM, standard error of the mean.

Crucially, cerebellar neuronal development is differentially affected across groups: MIA-exposed groups show transient effects on PC proliferation and arrested granule cell precursors (GCPs) in the EGL, while hypoxia-exposed groups show disruption of the postmigratory mature GC population in the IGL.

We next tested how these insult-specific phenotypes are reflected in the heterogeneous human preterm cerebellum by evaluating postmortem cerebellar tissue from preterm and term neonates (Fig. 2G), with documented major clinical inflammatory and hypoxic exposures (Supplementary Table 2-5). The human preterm EGL was persistently expanded, with elevated granule cell progenitor counts relative to term (Fig. 2H-I), recapitulating the MIA-exposed phenotype (Fig. 2D). In the postmigratory compartment, the preterm IGL surface area was reduced (Fig. 2J), though granule cell density within it remained comparable to term (Fig. 2K).

Following neuropathological analysis, we examined each insult-specific mouse gene program against the ranked human preterm versus term transcriptome, identifying the genes that mark this state and the signature that reproduces it. The isolated MIA and hypoxia mouse signatures showed non-significant concordance with the human (Supplementary Fig. 1A). By contrast, the sequential LPS-Hx program was the sole signature to be significantly enriched in the human preterm molecular state, with strong directional agreement (Supplementary Fig. 1A-B). This shared transcriptional signature was characterized by coordinated downregulation of programs governing neuronal communication, cell-cell signaling, and cerebellar granular layer formation (Supplementary Fig. 1C).

Together, these data support a model in which prematurity-related insults act on distinct stages of cerebellar development in an insult-specific manner, with their sequential combination reproducing the niche-specific structural and transcriptional features of the human preterm cerebellum.

### MIA- and hypoxia-driven spatial molecular programs shape the adult cerebellar transcriptome

To define the molecular programs underlying the niche-specific pathology identified above, we profiled the adult cerebellum, when behavioral deficits are fully manifested, using Visium spatial transcriptomics. This approach enabled spatial resolution of insult-specific transcriptional programs within the intact cerebellar architecture, linking developmental pathology to long-term circuit dysfunction.

By applying Coordinated Gene Activity in Pattern Sets (CoGAPS), an unsupervised factorization framework (Fig. 3A; Supplementary Fig. 2), we decomposed the spatial cerebellar transcriptome into 13 biologically distinct M (mouse)-patterns representing cell- (Supplementary Fig. 3) and insult-specific (Fig. 3B) gene expression modules.

**Fig. 3.**
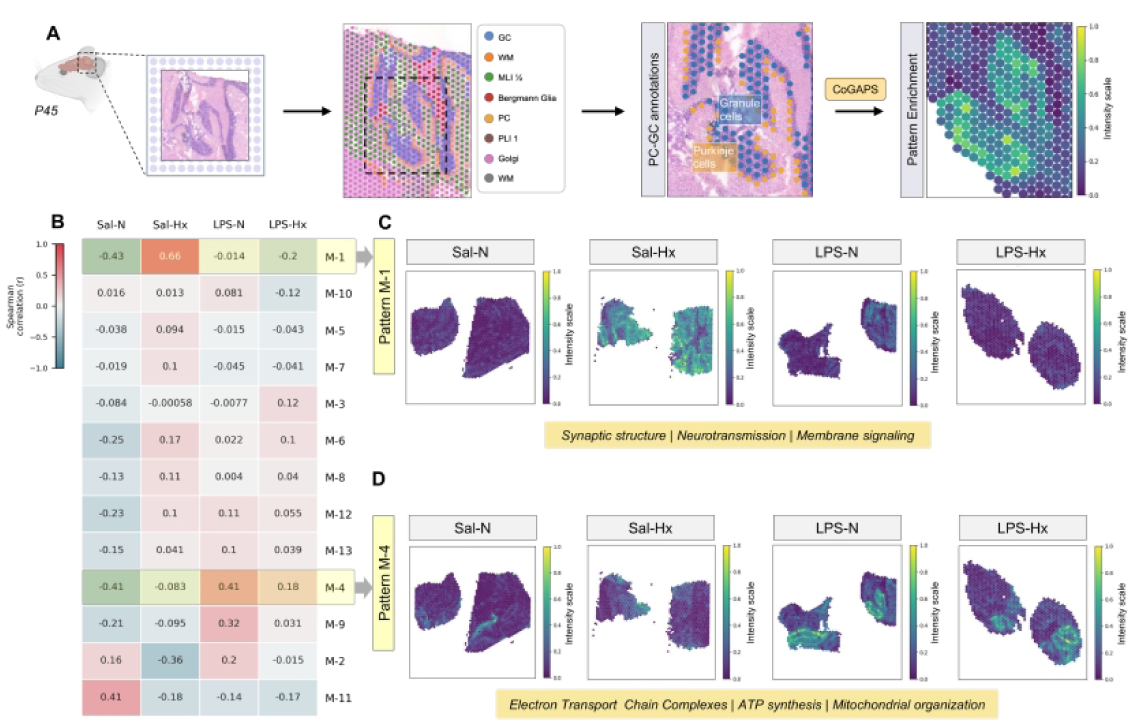
Insult-Specific Reprogramming of the Cerebellar Transcriptome. **(A)** Schematic illustration of the analytical pipeline for spatial transcriptomics and CoGAPS pattern discovery in mouse cerebellum. Hematoxylin and eosin-stained cerebellar tissue sections were processed for Visium spatial transcriptomics. All spatial transcriptomic samples were derived from P45 adult mice. Representative images demonstrate first cell-type annotation of cerebellar regions at 20x magnification and second, granule cells (blue) and Purkinje cells (orange) annotations at 40x magnification. Following CoGAPS analysis, gene expression patterns were mapped onto tissue architecture, with expression levels displayed on an intensity scale ranging from low (blue) to high (yellow). **(B)** Heatmap displaying Spearman correlation coefficients between CoGAPS-derived expression patterns and experimental conditions across all four groups. Correlation values range from high positive (red, *r* = 1) to high negative (blue, *r* = −1). **(C-D)** Spatial visualization of two representative CoGAPS patterns across experimental groups. Pattern M-1 (C) demonstrates enrichment for genes associated with synaptic structure, neurotransmission, and membrane signaling pathways. Pattern M-4 (D) demonstrates enrichment for genes associated with electron transport chain complexes, ATP synthesis, and mitochondrial organization. Expression intensity scale ranges from low (blue) to high (yellow). For all spatial transcriptomics analyses, *n* = 2 per group. Abbreviations: CoGAPS, Coordinated Gene Activity in Pattern Sets; GC, granule cell; Hx, hypoxia; LPS, lipopolysaccharide; M, mouse; MIA, maternal immune activation; MLI, molecular layer interneuron; N, normoxia; PC, Purkinje cell; PLI, Purkinje layer interneuron; Sal, saline; WM, white matter.

We focused on the M-patterns most strongly associated with each insult to define how the exposures engage in coordinated cell-type- and spatially organized transcriptional programs. Patterns M-1 and M-4 were the dominant patterns distinguishing insult exposures (Fig. 3B-D). M-1, strongly associated with hypoxia exposure (Fig. 3B-C), represented a generalized cerebellar transcriptional program (Supplementary Fig. 3) and comprised gene programs governing synaptic structure, vesicle trafficking, and neurotransmission (Supplementary Fig. 4A). In contrast, M-4, induced by MIA but not retained under sequential exposure (Fig. 3B, D), localized predominantly to Purkinje cells (Supplementary Fig. 3) and was defined by metabolic and mitochondrial gene programs (Supplementary Fig. 4B).

Together, these spatial programs indicate that hypoxia and MIA establish distinct, lasting molecular states within the adult cerebellum. Hypoxia alone was associated with a broad synaptic signature, whereas MIA induced a Purkinje-cell–associated metabolic program that was attenuated, rather than reinforced, by subsequent hypoxia. This pattern supports a priming-state model in which prior inflammation establishes a latent molecular vulnerability that is subsequently remodeled by hypoxic exposure.

### MIA primes the cerebellum for hypoxia-driven, progressive mitochondrial failure

MIA induced a cerebellar transcriptional program (pattern M-4) enriched for regulators of oxidative metabolism and mitochondrial control (Fig. 3B, D; Supplementary Fig. 4B), implicating the mitochondrion as a target of prenatal immune activation. We therefore asked whether this transcriptional signature was accompanied by a durable alteration in cerebellar bioenergetic state rather than a transient stress response.

Across developmental stages, the insults diverged sharply. Hypoxia alone produced early structural mitochondrial changes that largely resolved by adulthood (Fig. 4G, H, N, O), while MIA alone did not significantly alter bioenergetic function or mitochondrial density at either age (Fig. 4A, B, E, F, I, J). In contrast, MIA followed by hypoxia converted an initially compensated state into a progressive energetic failure, established by P45 (Fig. 4A-C, E-J, K-L, N-O). Thus, what begins as a subtle metabolic disturbance after sequential exposure evolves into a progressive collapse of cerebellar energy support.

**Fig. 4.**
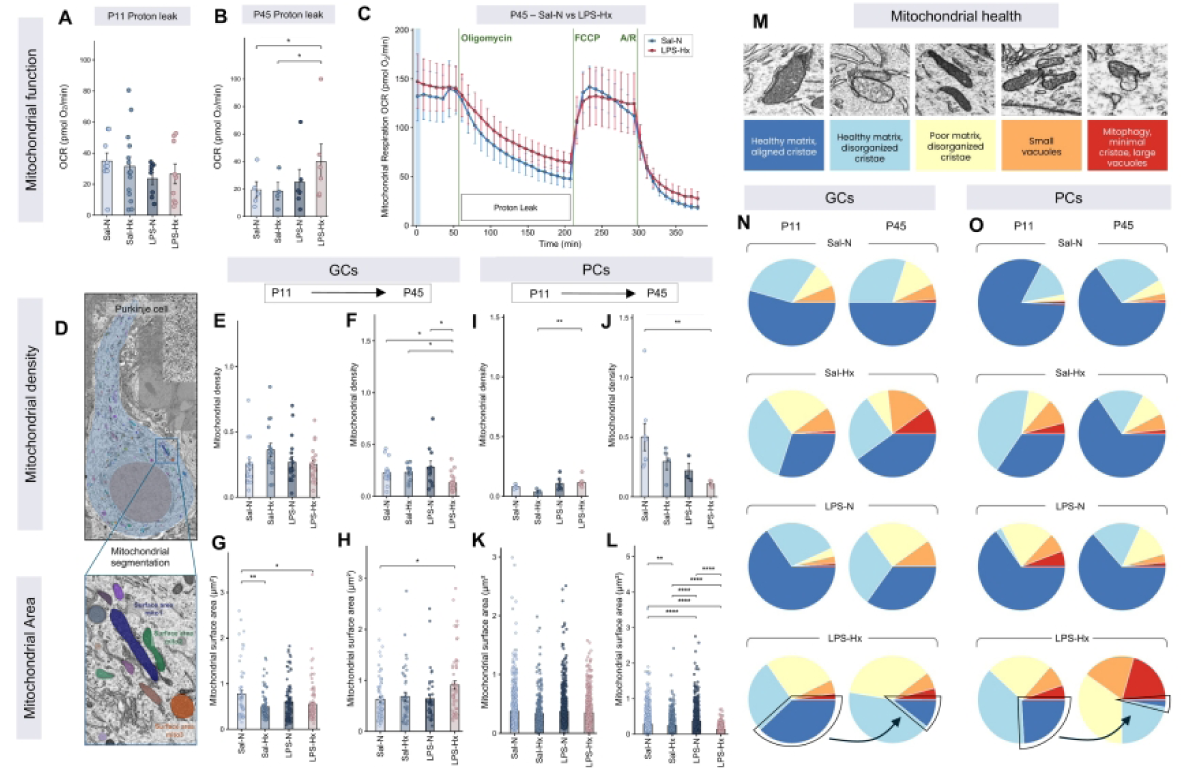
MIA-Induced Metabolic Priming Increases Vulnerability to Neonatal Hypoxia. (A–B) Quantification of proton leak rates at P11 (A) and P45 (B) (LPS-Hx versus Sal-Hx, \**P* = 0.031; LPS-Hx versus Sal-N, \**P* = 0.046) across experimental groups. **(C)** Representative Seahorse XF mitochondrial respiration oxygen consumption rate traces (µmol O₂/min) from cerebellum at P45, comparing Sal-N and LPS-Hx groups and displaying mitochondrial respiration across metabolic phases following sequential machine-applied injection of oligomycin, FCCP, and antimycin A/rotenone (A/R). The proton leak measurement window is indicated. **(D)** Schematic illustration of electron microscopy-based mitochondrial analysis in Purkinje cells. Top panel demonstrates mitochondrial identification and segmentation within a Purkinje cell soma; bottom panel displays representative examples of individual mitochondria with corresponding surface area measurements. **(E–H)** Mitochondrial morphometry in granule cells. (E–F) Mitochondrial density was assessed at P11 (E) and P45 (F) (Sal-N versus LPS-Hx, \**P* = 0.012; Sal-Hx versus LPS-Hx, \**P* = 0.010 LPS-N versus LPS-Hx, \**P* = 0.017). (G–H) Mitochondrial surface area (µm²) was quantified at P11 (G) (Sal-N versus Sal-Hx, \*\**P* = 0.0030; Sal-N versus LPS-Hx, \**P* = 0.023) and P45 (H) (Sal-N versus LPS-Hx, \**P* = 0.011). **(I–L)** Mitochondrial morphometry in Purkinje cells. (I–J) Mitochondrial density was assessed at P11 (I) (Sal-Hx versus LPS-Hx, \*\**P* = 0.0073) and P45 (J) (Sal-N versus LPS-Hx, \*\**P* = 0.0052). (K–L) Mitochondrial surface area (µm²) was quantified at P11 (K) and P45 (L)(Sal-N versus Sal-Hx, \*\**P* = 0.0087; Sal-N versus LPS-N, \*\*\*\**P* < 0.0001; Sal-N versus LPS-Hx, \*\*\*\**P* < 0.0001; Sal-Hx versus LPS-N, \*\*\*\**P* < 0.0001; Sal-Hx versus LPS-Hx, \*\*\*\**P* < 0.0001; LPS-N versus LPS-Hx, \*\*\*\**P* < 0.0001). **(M)** Representative electron micrographs illustrating mitochondrial ultrastructural quality assessment. A severity scoring system was applied: 1 (dark blue), healthy matrix with aligned cristae; 2 (light blue), healthy matrix with disorganized cristae; 3 (yellow), poor matrix with disorganized cristae; 4 (orange), small vacuoles; 5 (red), mitophagy with minimal cristae and large vacuoles. **(N–O)** Pie chart visualization of mitochondrial health distribution in granule cells (N) and Purkinje cells (O) across experimental groups at P11 and P45. Note the marked reduction in the proportion of healthy mitochondria (dark blue) from P11 to P45 in both Purkinje and granule cells of the LPS-Hx group. For (A), *n* = 8–14 per group. For (B), *n* = 5– 6 per group; one-way ANOVA with uncorrected Fisher’s LSD. For (C), *n* = 6 per group; one-way ANOVA with Tukey’s post hoc correction. For (E), *n* = 16–21 cells per group. For (F), n = 11–21 per group. For (E-F) Kruskal–Wallis test with Dunn’s post hoc test (Bonferroni-adjusted). For (G), *n* = 57–114 per group; one-way ANOVA with Tukey’s post hoc correction. For (H), *n* = 41–77 per group; one-way ANOVA with Tukey’s post hoc correction. For (I), *n* = 4–8 per group. For (J), *n* = 3–8 per group. For (I-J), Kruskal–Wallis test with Dunn’s post hoc test (Bonferroni-adjusted). For (K), *n* = 306–757 per group; one-way ANOVA with Tukey’s post hoc correction. For (L), *n* = 528–1783 per group; one-way ANOVA with Tukey’s post hoc correction. Data are presented as mean ± SEM; \**P* < 0.05, \*\**P* < 0.01, \*\*\*\**P* < 0.0001. Abbreviations: A/R, antimycin A/rotenone; EM, electron microscopy; FCCP, carbonyl cyanide-p-trifluoromethoxyphenylhydrazone; GC, granule cell; Hx, hypoxia; LPS, lipopolysaccharide; N, normoxia; OCR, oxygen consumption rate; PC, Purkinje cell; Sal, saline; SEM, standard error of the mean.

At P11, cerebellar functional bioenergetics (Seahorse OCR) were preserved across groups (Fig. 4A), yet by adulthood (P45) LPS-Hx animals showed elevated proton leak (Fig. 4B) on a background of preserved basal respiration (Fig. 4C), consistent with altered mitochondrial coupling and bioenergetic stress. Electron microscopy revealed the structural correlate of this failure: while neonatal GCs and PCs retained largely intact mitochondria, adult LPS-Hx cerebella displayed reduced mitochondrial density and altered surface-area distributions in both cell types (Fig. 4E–L), reflecting impaired mitochondrial turnover, a pattern we next examined directly by ultrastructural scoring.

Qualitative analysis^59^ confirmed a progressive accumulation of swollen, vacuolated mitochondria with severely disorganized cristae in the LPS-Hx group, most pronounced in Purkinje cells by P45. By contrast, hypoxia-only mice did not show this progressive decline: their overall mitochondrial-health distribution remained largely stable between P11 and P45, without the marked expansion of the mitophagic, large-vacuole fraction seen in LPS-Hx. This severe, end-stage fraction likewise remained minimal in MIA-only mice (Fig. 4M-O).

These findings dissociate three distinct metabolic trajectories of cerebellar injury: hypoxia alone produced a transient metabolic perturbation that resolved over development, MIA alone induced the M-4 metabolic program without a corresponding loss of mitochondrial function or density, and sequential exposure, without acutely altering mitochondrial function, established a latent susceptibility that progressed to mitochondrial failure over the postnatal period. This dissociation indicates that the LPS-Hx deficit reflects hypoxia acting on a substrate already primed by MIA, rather than the accumulated effects of the two exposures. Prior maternal immune activation therefore shifts the cerebellar response to subsequent hypoxia from a recoverable metabolic perturbation to a sustained energy-metabolic deficit.

### Insult-specific cell cycle arrest is linked with presynaptic-postsynaptic circuit mismatch

Persistent mitochondrial dysfunction engages checkpoint pathways that regulate neuronal proliferation and differentiation^60^. This raised the question of whether neonatal insult-induced energy failure leaves a durable molecular imprint on cell-cycle machinery within the structurally mature cerebellum, reflecting the GC and PC maturation phenotypes shown earlier (Fig. 2B-F). To capture these latent transcriptomic cues across distinct cerebellar layers, we applied Visium-based Tricycle analysis at P45 (Fig. 5A). Because adult granule cells are largely post-mitotic, we reasoned that differences in cell-cycle scores would reflect a persistent transcriptional signature of prior developmental arrest rather than ongoing proliferative activity.

**Fig. 5.**
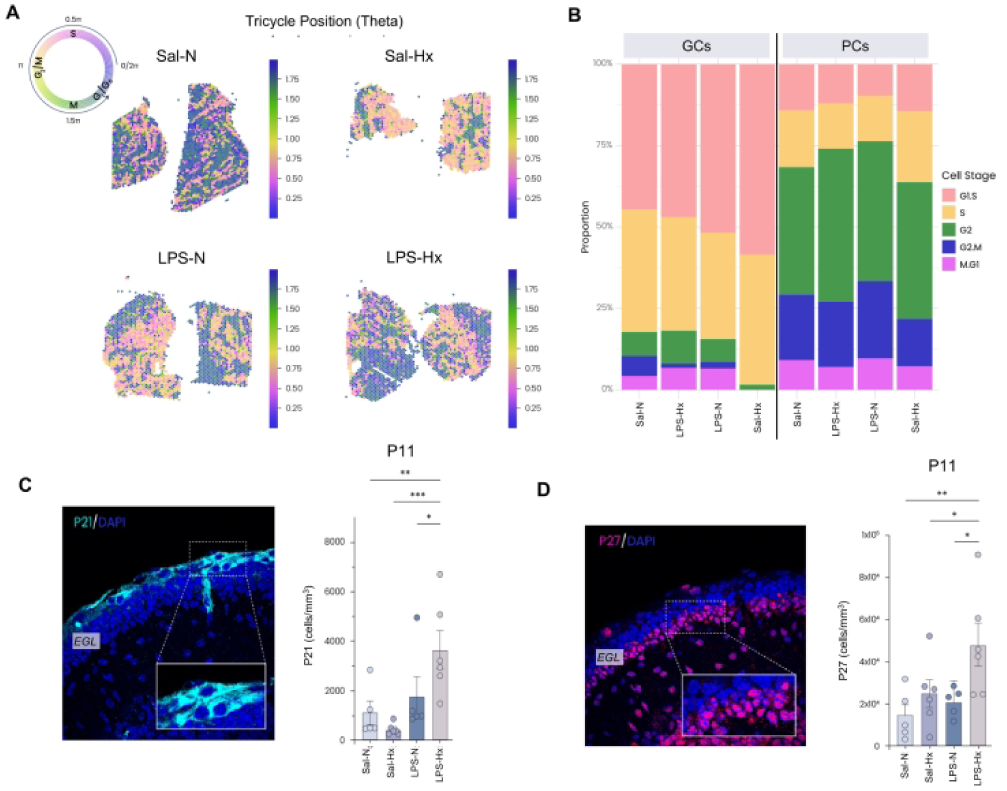
MIA Differentially Reconfigures Hx-Induced Alterations in Neuronal Cell Cycle Dynamics. **(A)** Spatial mapping of cell cycle phase distribution in mouse cerebellum. Top left schematic illustrates the cell cycle phases corresponding to the theta value with associated color scale: G1/G0 phase (blue/purple, θ < 0.25 and θ > 1.75), S phase (pink to yellow, θ = 0.5-1.0), and G2/M phase (green, θ = 1.25-1.75). Representative spatial transcriptomics images display cell-cycle phase distribution across cerebellar tissue for each experimental group. **(B)** Quantification of cell cycle phase proportions in granule cells and Purkinje cells across experimental groups. Stacked bar graphs display the percentage of cells in each cell-cycle phase, with colors corresponding to phases indicated in the legend. **(C)** Immunofluorescent detection of the cell-cycle inhibitor P21 in mouse cerebellar tissue. Left panel shows representative 60x magnification image of P21 (cyan) and DAPI (blue) staining from an LPS-Hx animal. Right panel displays quantification of P21-positive cells expressed as cells per cubic millimeter (Sal-N versus LPS-Hx, \*\**P* = 0.0070; LPS-N versus LPS-Hx, \**P* = 0.039; Sal-Hx versus LPS-Hx, \*\*\**P* = 8.0 × 10⁻⁴). **(D)** Immunofluorescent detection of the cell cycle inhibitor P27 in mouse cerebellar tissue (P11 mice). Left panel shows representative 60× magnification image of P27 (magenta) and DAPI (blue) staining from an LPS-Hx animal. Right panel displays quantification of P27-positive cells expressed as cells per cubic millimeter (Sal-N versus LPS-Hx, \*\**P* = 0.0042; LPS-N versus LPS-Hx, \**P* = 0.016; Sal-Hx versus LPS-Hx, \**P* = 0.029). For (C) and (D), *n* = 5-6 per group; one-way ANOVA with uncorrected Fisher’s LSD. Data are presented as mean ± SEM; \**P* < 0.05, \*\**P* < 0.01, \*\*\**P* < 0.001. Abbreviations: DAPI, 4′,6-diamidino-2-phenylindole; GC, granule cell; Hx, hypoxia; LPS, lipopolysaccharide; LSD, least significant difference; N, normoxia; PC, Purkinje cell; Sal, saline; SEM, standard error of the mean.

In granule cells, neonatal hypoxia alone (Sal-Hx) was associated with a coordinated depletion of the M/G1, G2/M, and G2 phases together with an accumulation of G1/S (Fig. 5B), suggesting a stalled progression through the cell cycle and arrested postmigratory granule cell maturation. In contrast, the MIA-exposed groups (LPS-N and LPS-Hx) showed selective depletion of the G2/M phase relative to Sal-N (Fig. 5B), consistent with impaired entry into mitosis. Purkinje cells showed no comparable insult-driven cell-cycle differences (Fig. 5B), indicating that their molecular vulnerability at this mature stage is independent of proliferative checkpoint activity.

Next, we demonstrated that these P45 cell-cycle abnormalities were already evident during early granule cell development by examining expression of P21 and P27, the principal cyclin-dependent kinase inhibitors enforcing cell-cycle arrest, at P11, when granule cell progenitors are actively proliferating within the EGL (Fig. 5C-D). Both inhibitors were markedly elevated in LPS-Hx cerebella compared with all other groups (Fig. 5C-D). This pattern indicates that sequential exposure engages checkpoint arrest throughout the GC maturational trajectory, linking neonatal cell-cycle disruption to the lasting cerebellar phenotypes observed in adulthood.

The persistent immature transcriptional state of granule cells and the hypoxia-associated synaptic program (M-1, Fig. 3B-C) pointed to cerebellar circuit assembly as the level at which these insults act. Because postmigratory granule cell maturation depends on activity-dependent and trophic cues from Purkinje cells^61,62^, we examined the two integrated arms of this circuit, VGLUT2⁺ climbing fiber input onto Purkinje cells and Purkinje cell dendritic length and complexity (Fig. 6A-F), alongside the GC↔PC interactome across groups (Supplementary Fig. 5A-D).

**Fig. 6.**
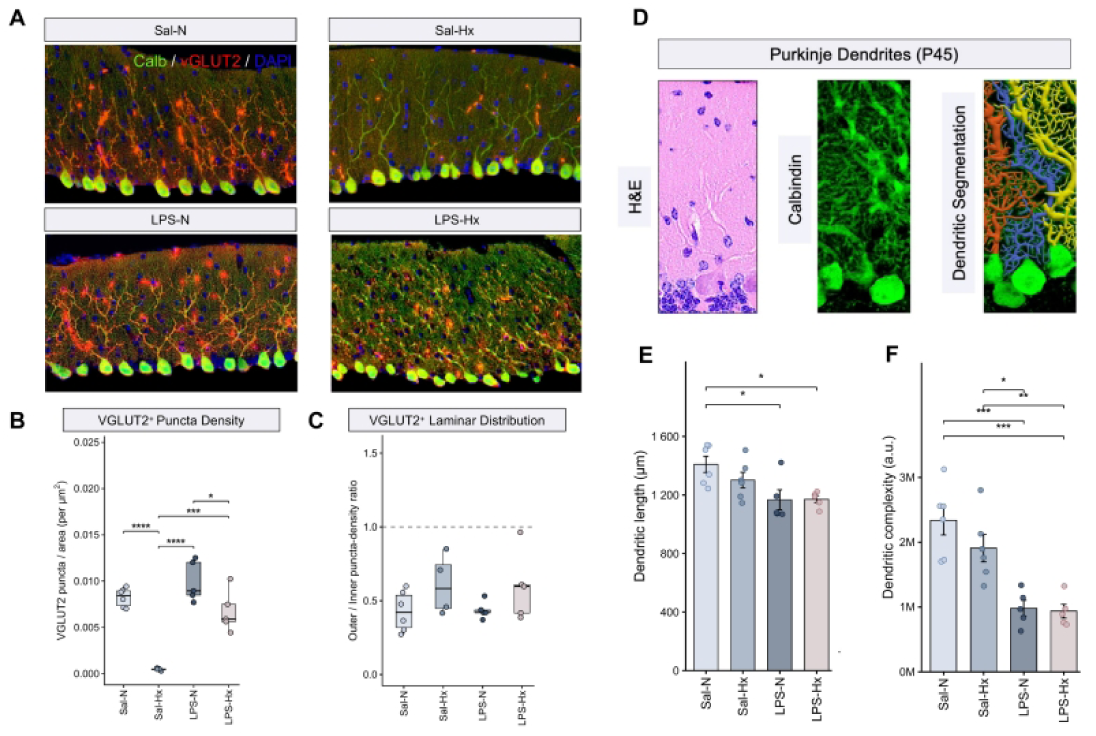
Perinatal Insults Rewire the Cerebellar Connectome through Divergent Disruption of Cell-Cell Signaling Networks. **(A)** Representative confocal images of P45 cerebellar tissue displaying calbindin-immunostained Purkinje cells and dendrites (green), VGLUT2-immunostained excitatory synaptic puncta (red), and DAPI nuclear counterstain (blue) across all four experimental groups (Sal-N, Sal-Hx, LPS-N, LPS-Hx) at 40x magnification. **(B)** Quantification of VGLUT2-positive puncta density expressed as VGLUT2 puncta per area (per µm²) (Sal-N versus Sal-Hx, \*\*\*\**P* < 0.0001; Sal-Hx versus LPS-N, \*\*\*\**P* < 0.0001; Sal-Hx versus LPS-Hx, \*\*\**P* = 2.0 × 10⁻⁴; LPS-N versus LPS-Hx, \**P* = 0.035). **(C)** VGLUT2-positive laminar distribution expressed as the ratio of outer to inner puncta density across experimental groups. Horizontal dashed line indicates a balanced ratio (ratio = 1.0). **(D)** Schematic illustration of the Purkinje cell dendritic arborization quantification pipeline at P45. Left panel displays hematoxylin and eosin-stained cerebellar section; middle panel shows representative calbindin-immunostained Purkinje cell imaged by confocal microscopy; right panel demonstrates Neurolucida dendritic segmentation. All images captured at 100x magnification. **(E)** Quantification of Purkinje cell dendritic length (µm) at P45 (Sal-N versus LPS-N, \**P* = 0.023; Sal-N versus LPS-Hx, \**P* = 0.026). **(F)** Quantification of Purkinje cell dendritic complexity (a.u.) at P45 (Sal-N versus LPS-N, \*\*\**P* = 3.0 × 10⁻⁴; Sal-N versus LPS-Hx, ***P = 2.0 × 10⁻⁴; Sal-Hx versus LPS-N, \**P* = 0.010; Sal-Hx versus LPS-Hx, \*\**P* **=** 0.0078). For (B), (E), and (F), *n* = 4–6 per group; one-way ANOVA with Tukey’s post hoc correction. For (C), *n* = 4–6 per group; Kruskal–Wallis test with Dunn’s post hoc test (Bonferroni-adjusted). Data are presented as mean ± SEM; \**P* < 0.05, \*\**P* < 0.01, \*\*\**P* < 0.001, \*\*\*\**P* < 0.0001. Abbreviations: a.u., arbitrary units; Calb, calbindin; DAPI, 4′,6-diamidino-2-phenylindole; GC, granule cell; H&E, hematoxylin and eosin; Hx, hypoxia; LPS, lipopolysaccharide; N, normoxia; PC, Purkinje cell; Sal, saline; SEM, standard error of the mean; VGLUT2, vesicular glutamate transporter 2.

The two insults disrupted opposite arms of this circuit. Hypoxia alone produced a presynaptic phenotype, with near-complete depletion of VGLUT2⁺ puncta onto Purkinje cells whose dendritic architecture remained largely preserved (Fig. 6B, E-F). Additionally, it redirected the baseline GC-PC interactome (Sal-N; Supplementary Fig. 5A) from programs supporting adhesion, axon guidance, and glutamatergic circuit assembly toward a stress- and injury-associated remodeling state, marked by APOE→LRP1, SPARC→FGFR1, and GDF10 signaling (Supplementary Fig. 5B).

Conversely, MIA produced a predominantly postsynaptic phenotype, reducing Purkinje cell dendritic length and complexity in both the LPS-N and LPS-Hx groups (Fig. 6E-F). In the sequential LPS-Hx group, prior MIA attenuated the presynaptic depletion observed with hypoxia alone, with VGLUT2⁺ input maintained above Sal-Hx levels and comparable to Sal-N controls (Fig. 6B), within an inner-molecular-layer-predominant distribution that was conserved across groups (Fig. 6C). However, this preservation of presynaptic input did not indicate physiological recovery. Instead, sequential exposure redirected the hypoxic response toward a mismatched circuit state, in which preserved afferent input converged onto a constricted and atrophied postsynaptic Purkinje cell territory.

Consistent with this structural divergence, MIA alone caused a limited reorganization of the adult GC–PC interactome, selectively rerouting guidance- and adhesion-associated signaling, including OMG→RTN4RL1 and L1CAM→CD9 (Supplementary Fig. 5C). By contrast, sequential LPS-Hx engaged degeneration- and matrix-remodeling pathways absent from either single insult, including APP→TNFRSF21, TNC→CNTN1, and ADAM10→TSPAN15 (Supplementary Fig. 5D), suggesting that sequential exposure produces a distinct remodeling state rather than a simple summation of MIA and hypoxia effects.

Together, these findings suggest that prematurity-associated insults impair GC–PC circuit maturation not through uniform synaptic loss, but through maladaptive reorganization of presynaptic input, Purkinje cell dendritic architecture, and intercellular signaling. This maladaptive remodeling links persistent granule cell immaturity to altered Purkinje cell afferent integration, identifying a potential cellular substrate for later socio-cognitive dysfunction.

### Sequential insults transform hypoxia-induced social and motor dysregulation

Prematurity-associated insults produced distinct disruptions in cerebellar circuit assembly and sensorimotor adaptation, pointing to impaired cerebellar integration. We therefore examined whether these circuit-level changes extended to higher-order behaviors, including social engagement. To capture social approach at the level of broad chamber-based behavior, we used the three-chamber assay to assess gross sociability (Fig. 7A–B, Supplementary Fig. 6A-B).

**Fig. 7.**
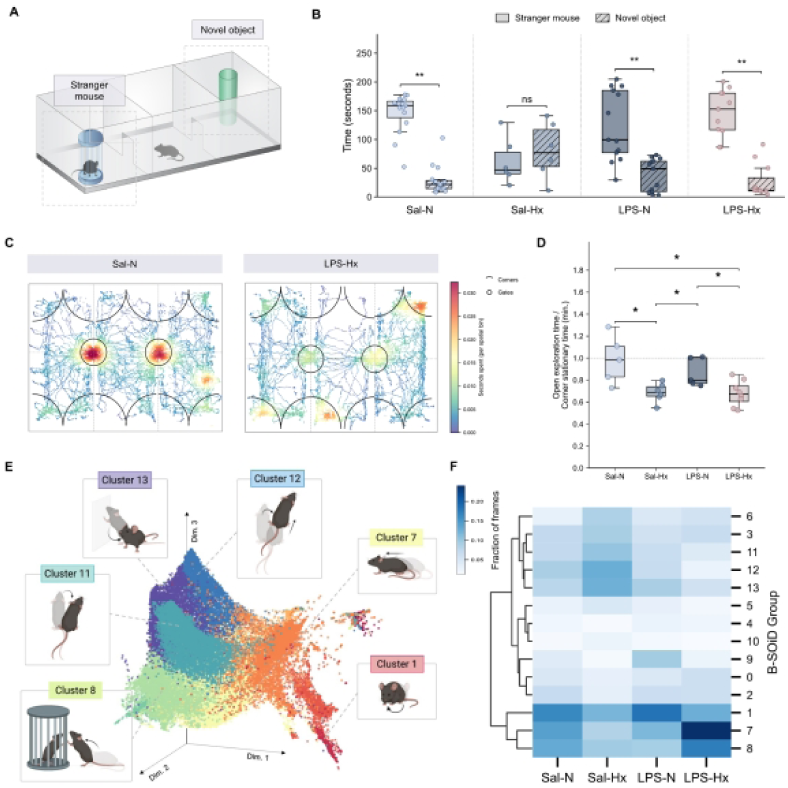
MIA and Hx Elicit Divergent Social, Exploratory, and Kinematic Phenotypes. **(A)** Schematic illustration of the three-chambered social approach assay used to evaluate sociability in mice. Left chamber contains an unfamiliar stimulus mouse housed within a cylindrical wire cage; middle chamber represents the arena where the test subject is placed; right chamber contains a novel object (green cylinder). **(B)** Quantification of time spent in the social chamber versus the object chamber across experimental groups. Plain boxplots represent time spent in the social chamber and striped boxplots represent time spent in the object chamber. Within-group comparisons: Sal-N, \*\**P* = 0.0044; Sal-Hx, ns; LPS-N, \*\**P* = 0.0068; LPS-Hx, \*\**P* = 0.0068. **(C)** Representative movement traces from individual Sal-N and LPS-Hx mice during the habituation phase (stimulus-free) assessment. Tracked trajectories are displayed with chamber corners demarcated by black semicircles, while gate areas are demarcated by black full circle. Vertical dashed light grey lines mark chamber boundaries and horizontal dashed grey line marks chamber midline. Heatmap scale indicates time spent per spatial bin ranging from low (blue) to high (red). **(D)** Quantification of exploratory behavior expressed as the ratio of open exploration time to corner stationary time across experimental groups (Sal-N versus Sal-Hx, \**P* = 0.035; Sal-N versus LPS-Hx, \**P* = 0.032; Sal-Hx versus LPS-N, \**P* = 0.033; LPS-N versus LPS-Hx, \**P* = 0.030). **(E)** Three-dimensional visualization of the 14 kinematic behavioral clusters identified by the B-SOiD algorithm. Representative illustrations depict kinematic motifs for selected clusters, including Cluster 1, Cluster 7, Cluster 8, Cluster 11, Cluster 12, and Cluster 13. **(F)** Heatmap displaying the frequency of kinematic motifs across experimental groups, expressed as fraction of video frames classified into each of the 14 behavioral clusters. Color intensity ranges from light blue (lower fraction, 0.05) to dark blue (higher fraction, 0.20). Hierarchical clustering dendrogram shows the frequency of the different behavioral clusters across the four experimental groups. For (B) and (E-F), *n* = 6–15 per group; within-group comparisons performed using paired Wilcoxon signed-rank test. For (D), *n* = 5-8 per group; pairwise two-sample Welch’s t-test across the four groups. Data are presented as mean ± SEM; \**P* < 0.05, \*\**P* < 0.01. Abbreviations: B-SOiD, Behavioral Segmentation of Open-field In DeepLabCut; Hx, hypoxia; LPS, lipopolysaccharide; N, normoxia; ns, not significant; Sal, saline; SEM, standard error of the mean.

This initial chamber-based readout revealed mixed social phenotypes across groups. Sal-N controls displayed the expected pattern of sociability in this assay, preferentially occupying the chamber containing a stranger conspecific over the non-social chamber. Isolated hypoxia (Sal-Hx) disrupted this pattern, resulting in reduced orientation toward the social cue (Fig. 7B). In contrast, MIA-exposed mice, both LPS-N and LPS-Hx, showed elevated social-chamber occupancy, which at first glance suggested preserved social preference (Fig. 7B).

Next, we used the stimulus-free habituation phase as an open-field context to assess whether prematurity-associated sensorimotor deficits emerged during spontaneous exploratory activity before social exposure (Fig. 7C-D). Hypoxia-exposed groups (Sal-Hx, LPS-Hx) showed reduced open-field exploration, increased immobility, and greater corner-associated occupancy, consistent with a constrained exploratory strategy (Fig. 7D). Despite comparable baseline exploratory restriction across hypoxia-exposed groups, the opposing social-chamber outcomes raise the question of how each group organizes its investigatory behavior within the social context.

Thus, we examined the kinematic components of social behavior to determine whether the contrasting patterns of sociability and exploratory activity reflected changes in motor structure and the organization of stimulus-rich investigation, by applying an unsupervised algorithm (B-SOiD) which segmented tracked movements into 14 unbiased kinematic motif clusters (Fig. 7E, Supplementary Table 8, Supplementary Fig. 7). Group-wise frequency analysis of kinematic motif clusters separated the four groups by motif usage, separating Sal-Hx and, most prominently, LPS-Hx from controls and isolated MIA (Fig. 7F).

Specifically, Sal-Hx mice showed a predominance of repetitive body turning and reorientation (clusters 11, 12, and 13) with reduced use of the interactive approach motifs (clusters 7 and 8; Fig. 7F), a stereotyped, low-interactivity movement pattern that was concentrated in the novel-object area (Supplementary Fig. 8; Supplementary Fig. 9). Isolated MIA (LPS-N) produced a distinct reorganization, reducing the same interactive motifs (clusters 7 and 8) while shifting toward low-amplitude orienting and sensory sampling (body and head turning, clusters 1 and 9; Fig. 7F), indicating low investigatory engagement despite an intact gross social preference. In contrast, LPS-Hx mice displayed a hyper-interactive pattern of social investigation, dominated by directed approach and active exploratory engagement (clusters 7 and 8; Fig. 7F) at frequencies exceeding controls, expressed preferentially in the stranger-mouse chamber (Supplementary Fig. 8; Supplementary Fig. 9).

Together, these findings demonstrate that prematurity-related insults drive distinct, insult-specific socio-motor strategies. While control and LPS-N mice maintained a normative social preference, Hx alone abolished this baseline sociability, inducing a sensory-disengaged kinematic profile linked to impaired sensorimotor adaptation. Conversely, sequential LPS-Hx exposure preserved overall social interest but altered interaction qualities, marked by hyperactive and repetitive motor motifs. Thus, early-life insults do not merely deplete social interest, as seen in Hx, but can instead selectively reorganize the qualitative execution of investigation by disrupting the fine coordination between sensory input, locomotor state, and active engagement.

## Discussion

By modeling the interplay of maternal immune activation and neonatal hypoxia, this study provides direct evidence that prematurity-associated insults drive divergent disruptions in cerebellar circuitry, thereby shaping long-term motor and social phenotypes. Prior work has established that MIA and hypoxic injury each contribute to neurodevelopmental disability through inflammatory, synaptic, and metabolic perturbations, primarily characterized in cortical and hippocampal circuits^63,64,65,66^. However, these insults frequently co-occur in preterm infants, raising the possibility that their consequences depend not only on the identity of the insult but also on developmental timing and sequence. The cerebellum, despite its rapid and protracted development, which renders it disproportionately vulnerable to perinatal injury^16^, has not been fully explored. By recapitulating key features of the human preterm cerebellum, this single- and sequential-insult model exposes sequence-specific vulnerabilities in synaptic organization and metabolic reserve.

A central insight is that MIA and hypoxia perturb cerebellar circuitry through distinct pathways, with prior MIA reshaping the cerebellar response to subsequent hypoxic exposure. Within the canonical cerebellar circuit, Purkinje cells integrate parallel fiber input from granule cells with climbing fiber input from the inferior olive^67,68^. Our findings suggest that MIA and Hx impose synapse-specific vulnerabilities within this circuit: isolated Hx disrupts the presynaptic afferent compartment, marked by VGLUT2 depletion, whereas MIA, alone or preceding Hx, preserves these glutamatergic terminals while constraining Purkinje cell dendritic complexity and axonal growth. We propose that afferent preservation in the sequential condition may reflect a shift in hypoxic vulnerability driven by inflammatory preconditioning and altered developmental timing. Consistent with prior work showing that low-grade inflammation can induce ischemic tolerance and reduce excitotoxic injury after subsequent hypoxic-ischemic exposure^69^, MIA may reduce the susceptibility of glutamatergic afferents to Hx-induced disruption. At the same time, delayed cerebellar maturation may move these terminals outside a period of peak metabolic and synaptogenic vulnerability^70^. Thus, sequential exposure spares the presynaptic VGLUT2 network but leaves it coupled to Purkinje cells with reduced integrative capacity, producing a preserved-yet-mismatched circuit architecture.

The structural mismatch produced by sequential exposure may represent a form of maladaptive plasticity that reframes the origins of prematurity-associated behavioral sequelae. Rather than producing discrete deficits through overt circuit loss, sequential insults may preserve cerebellar circuitry in an altered configuration, giving rise to graded and variable behavioral outcomes. In this surviving but miswired state, disrupted circuit organization may provide a mechanistic basis for the overlapping social and motor features of the preterm behavioral phenotype, including autism spectrum disorder and developmental coordination disorder^71^.

The cerebellum generates predictive estimates that coordinate the timing and sequencing of action^72^. Preserved climbing-fiber drive onto a constricted Purkinje cell dendritic arbor may distort, rather than silence, cerebellar computation. Afferent excitation remains intact but is processed through a compromised integrative compartment, creating an excitation-integration mismatch that degrades predictive motor sequencing. This model explains how social motivation can be preserved even as the sensorimotor organization required for social approach becomes disordered, linking social and motor impairment after preterm birth to the preservation of maladaptive circuits rather than afferent loss.

Beyond altered connectivity, our findings identify progressive mitochondrial failure as another major mechanism linking early-life insults to lasting cerebellar dysfunction. In the sequential exposure group, early metabolic compensation was followed by delayed bioenergetic collapse, indicating that MIA establishes a latent mitochondrial vulnerability that is unmasked by later hypoxic stress. This timing is mechanistically important: rather than depleting metabolic capacity outright, MIA constrains cerebellar bioenergetic reserve, so that failure emerges when the increasing energetic demands of postnatal circuit maturation, or a superimposed hypoxic challenge, exceed adaptive capacity. Proliferating neonatal progenitors significantly rely on aerobic glycolysis, so a constraint on mitochondrial expansion remains bioenergetically silent^73,74^ and surfaces only as post-mitotic maturation shifts the tissue toward oxidative phosphorylation and raises oxidative demand^73^. This positions inflammatory priming as a temporally extended window of vulnerability rather than an immediate metabolic disruption, offering a potential explanation for the prolonged susceptibility observed in preterm infants beyond the immediate perinatal period^6,14^.

Together with the synaptic and circuit changes, these data indicate that early-life insults shape long-term cerebellar function through interacting mechanisms of afferent disruption, impaired Purkinje cell integration, delayed granule cell maturation, and constrained mitochondrial adaptation. The identification of a primed mitochondrial state further suggests opportunities for targeted metabolic strategies^75^ and positions MIA as a developmental modifier of hypoxic injury rather than a parallel insult acting independently. In this framework, the same hypoxic exposure may have different consequences depending on the maturational and metabolic state established by prior prenatal inflammation. This has translational relevance, as risk stratification based only on the presence or severity of individual insults may miss the biological importance of sequence, timing, and developmental state. These findings also suggest that interventions aimed at preserving cerebellar function may need to consider not only molecular protection and bioenergetic support, but also the timing of locomotor-dependent circuit refinement during early development, including through existing NICU practices such as physical therapy and developmentally guided sensorimotor care.

While our study modeled these insults on a neurotypical developing brain, clinical preterm birth is driven by diverse maternal, placental, and fetal etiologies that may independently compromise neural integrity, obscuring the effects of MIA and hypoxia in ways that cannot be fully replicated experimentally. The full etiological heterogeneity of prematurity inherently exceeds what any single model can capture. The present work focused exclusively on the cerebellum; the contribution of other brain regions to the observed behavioral phenotypes was not assessed. Postnatal survival heterogeneity within the human comparator cohort can influence comparability across the developmental window assessed. Both sexes were represented, though not in balanced proportions, so sex-specific effects were not analyzed separately. Despite these limitations, the controlled sequencing of insults within a conserved cerebellar architecture provides mechanistic resolution inaccessible in human studies, linking specific cellular disruptions to functional outcomes.

Collectively, these findings move prematurity-associated cerebellar injury from a model of nonspecific vulnerability toward a mechanistic framework in which the timing and sequence of perinatal insults define cerebellar development and outcomes. By linking developmental neuropathology to spatial molecular programs, mitochondrial state, circuit organization, and adult behavior, this work shows that prematurity-associated cerebellar injury is not a single endpoint, but a set of developmentally programmed trajectories shaped by exposure history. This framework supports precise biomarkers, precise behavioral phenotyping, and targets for designing interventions aimed at preserving cerebellar developmental resilience in high-risk preterm infants.

## Supporting information

Supplemental material for manuscript

## Data availability

All data, code, and scripts generated and used in this study are publicly available at https://github.com/kratimenoslab/Mouse-Prematurity-Insults. Any additional information required to reanalyse the data reported in this paper is available from the corresponding author upon request.

## Funding

This work was supported by the Raynor Cerebellum Project (P.K.); the National Institutes of Health (R37NS109478, Javits Award, V.G.; K12HD001399 Child Health Research Career Development Award, P.K.; R21NS135088, I.K.; and P50HD105328, District of Columbia Intellectual and Developmental Disabilities Research Center award, V.G., from the Eunice Kennedy Shriver National Institute of Child Health and Human Development); the American SIDS Institute (P.K.); and a Children’s National Board of Visitors’ Grant (P.K.).

## Competing interests

The authors report no competing interests.

## Supplementary material

Supplementary material accompanies this manuscript.

## Notes

https://github.com/SteinOBrienLab/MIA_CB_Visium/tree/main

### Competing Interest Statement

The authors have declared no competing interest.

### Summary of Updates

This revised version of the manuscript includes updated text throughout the Results and Discussion sections to improve clarity, flow, and interpretation of the findings. We incorporated new LIANA cell-cell communication analyses to better define condition-specific signaling changes within the developing cerebellum and to support the proposed mechanisms linking early-life insults to altered cerebellar circuitry. The manuscript was also updated to include new and revised social behavior results, with clearer interpretation of group-specific differences in chamber occupancy and social cue engagement.

