## Supplemental material for manuscript for "Prematurity insults remodel cerebellar development and behavior"

### Materials and methods

#### Ethics

This research was conducted in accordance with the ethical guidelines and approval of the Institutional Review Board (IRB) at Children's National Research Institute. All human samples included in this study were voluntarily donated with informed consent and permission to use provided by patients' families under IRB-approved protocols (IRB #00011850, IRB #00015350), which cover sample acquisition, clinical data abstraction, analysis, and data sharing. Source of tissue sections are Children's National Hospital Department of Pathology and NIH NeuroBioBank.

#### Prematurity-associated insults mouse model

Wildtype C57BL/6 (Jackson Laboratory, strain #000664) and CD1 [Charles River Crl:CD1(ICR)] mice were maintained in the animal facility of the Children's National Hospital. All mouse procedures were approved by the Institutional Animal Care and Use Committee (IACUC #00030473) of Children's National Hospital, conducted in accordance with the Guide for the Care and Use of Laboratory Animals (National Institute of Health). Mice were maintained in the animal facility under a 12-hour dark–light cycle, constant temperature (20–26 °C) and humidity maintenance (40–60%), and access to food and water ad libitum.

Group allocation was randomized at two levels. Timed-pregnant C57BL/6 dams (vaginal plug = embryonic day 0.5) were randomly assigned to the saline or lipopolysaccharide (LPS) condition in a 1:1 ratio using a random-number generator (R), with the allocation sequence generated by an investigator not involved in outcome assessment. At postnatal day 2, litters were randomly assigned to normoxia or hypoxia using the same procedure; pups assigned to hypoxia were pooled and cross-fostered to CD1 dams and the foster litter was placed in the hypoxia chamber (see *Maternal care and cross-fostering*), whereas normoxia pups remained with their biological C57BL/6 dam. This yielded the four experimental groups:

1. Saline-Normoxia (Sal-N): pregnant C57BL/6 dams were injected with normal saline at embryonic day (E) 15 and E16 and the pups were born and maintained under normal oxygen concentration conditions (N).
2. Saline-Hypoxia (Sal-Hx): pregnant C57BL/6 dams were injected with normal saline at E15 and E16 and the pups were postnatally exposed to hypoxia (Hx; with foster dams) from postnatal day (P) 3 to P11.
3. LPS-Normoxia (LPS-N): pregnant C57BL/6 dams were injected with LPS (Millipore Sigma L6529, lot: 0000160286) at E15 and E16 and maintained in N.

4. LPS-Hypoxia (LPS-Hx): pregnant C57BL/6 dams were injected with LPS (Millipore Sigma L6529, lot: 0000160286) at E15 and E16 and the pups were exposed to Hx (with CD1 foster dams) from P3 to P11.

Each experimental group comprised pups derived from at least six independent litters, and littermates from a given dam were distributed across, rather than concentrated within, the groups compared. Although the hypoxia condition was assigned at the litter level, the individual animal served as the experimental unit for all analyses, and no group was composed of pups from a single litter. Because pups from multiple litters contributed to each group, litter was treated as a source of biological variability; the distribution of litters across groups minimized litter-driven confounding, and no single litter dominated any group.

**Maternal immune activation** Pregnant C57BL/6 dams received two intraperitoneal injections of LPS at E15 and E16, corresponding to 22–24 weeks of human gestation, the gestational window of peak preterm birth incidence. LPS was diluted in 0.9% normal saline to a final concentration of 33 μg/mL and injected at 4.5 μL/g of mouse weight. LPS activates Toll-like receptor 4 (TLR4)[19], a key innate immune mediator at the maternal-fetal interface that responds to both bacterial endotoxin and endogenous danger-associated molecular patterns, initiating a pro-inflammatory cascade that promotes uterine contractility and cervical remodeling[20,21,22]. This recapitulates the inflammatory signaling pathway through which clinical chorioamnionitis drives spontaneous preterm labor[19,23]. In our colony, LPS administration consistently shortened gestation to E18–E19 compared to the typical term of ~E20, reliably triggering spontaneous preterm birth.

**Hypoxia** The hypoxic groups were placed inside a sealed, automated hypoxia chamber (BioSpherix, Redfield, NY) in which FiO2 was maintained at 10.5–11% by nitrogen displacement, with continuous O2 monitoring, CO2 absorption, and chamber temperature/humidity matched to housing. The chamber was opened only briefly (<5 minutes) for scheduled husbandry, minimizing environmental fluctuations. Normoxia groups were housed under identical conditions at 21% O2. Hypoxia onset at P3 was selected because earlier exposure (P1–P2) is precluded by high neonatal mortality due to cardiovascular and thermoregulatory immaturity, and the P3–P11 exposure window follows the established chronic sublethal hypoxia paradigm validated by our group[14] for modeling neonatal brain injury. This window also recapitulates the sustained cerebral hypoxia that preterm infants experience during early postnatal life due to cardiorespiratory immaturity and impaired cerebral autoregulation[24].

**Maternal care and cross-fostering** Pups in the Sal-Hx and LPS-Hx groups were cross-fostered with CD1 dams at P2; Sal-N and LPS-N pups remained with their biological C57BL/6 dams. The foster CD1 dam always remained inside the chamber with the litter to ensure uninterrupted maternal care. Following completion of the hypoxic exposure at P11, pups in the hypoxic groups underwent a brief reacclimatization to normoxic conditions and were then returned to standard laboratory housing, where they remained until weaning at P21[14]. Cross-fostering with CD1 dams is standard practice for chronic neonatal hypoxia paradigms, as C57BL/6 dams exhibit inadequate maternal behavior under prolonged hypoxic conditions, resulting in poor pup survival and inconsistent nurturing[25,26]. CD1 dams were selected because they exhibit robust maternal behavior, large litters, and strong stress tolerance, features that support optimal neonatal survival during prolonged hypoxic exposure and reduce variability caused by inadequate maternal care. Both male and female pups were included in all experimental groups (Sal-N: male = 15, female = 3; Sal-Hx: male = 3, female = 5; LPS-N: male = 12, female = 2; LPS-Hx: male = 6, female = 4), but the distribution was not balanced enough to permit meaningful sex-specific statistical analyses. Because the study was not powered to assess sex-by-factor interactions, all results were analyzed at the group level, and sex is acknowledged as a potential source of variability.

#### Mouse and human histology and immunostaining

Histological analysis was performed at P11 and P45 in mice from all condition groups (Sal-N, Sal-Hx, LPS-N, and LPS-Hx; $n$ = 4-5 mice per group) and immunofluorescent analysis was performed again in mice from all condition groups at both P11 and P45 timepoints ($n$ = 4-5 mice per group at each time point). P11 was selected as the acute post-insult, developmentally relevant neonatal stage (external granule cell layer or EGL present, active GC migration, Purkinje cell or PC arborization), while P45 served as the early adulthood timepoint to assess long-term persistence of these effects. Tissue collection was performed within ~24 hours of the end of hypoxic exposure for P11 samples and within ~24 hours of the final behavioral assessment for P45 samples, ensuring that measurements reflected stable effects rather than acute activity-dependent or exposure-dependent changes.

**Histological and immunofluorescence experimental procedures** Mice were anesthetized with isoflurane and perfused transcardially with 0.1 M phosphate buffered saline (PBS), pH 7.4, followed by 4% paraformaldehyde (PFA). Brains were postfixed in 4% PFA overnight. Then, brains were placed in sucrose solution (15% sucrose, 85% PBS) overnight, and again in new sucrose solution (30% sucrose, 70% PBS) overnight. Following sucrose equilibration, brains were cut midsagittally, embedded in optimal cutting temperature (OCT) compound, and stored at -80 °C to preserve RNA integrity and antigenicity for downstream molecular and histological analyses. Prior to immunofluorescence, brains were sectioned on a cryostat at 40 μm thickness. For each animal, 3-4 midsagittal vermal sections were collected per endpoint. A subset of sections was mounted on slides and stained with hematoxylin and eosin (H&E) using standard neurohistological procedures, while the remaining sections were transferred into cryoprotectant solution (30% glycerol, 30% ethylene glycol, 40% PBS) and stored at 4 °C until staining. All quantitative analyses were performed on midsagittal whole-section images acquired at 20× magnification using QuPath software, with the animal, rather than individual sections or fields, serving as the biological unit for statistical analysis. Immunofluorescence analysis was performed on free-floating sections, which were blocked for two hours in PBST (0.3% TritonX-100), 20% normal donkey serum (NDS), and 1% bovine serum albumin (BSA); followed by overnight incubation at $4^{0}$C in primary antibodies diluted in PBST and 1% NDS, and 1% BSA. The brain sections were incubated with species appropriate secondary antibodies for two hours at room temperature. Information on primary and secondary antibody and dilutions can be found in Supplementary Table 1. The brain sections were mounted with DAPI (Vector Laboratories, USA) on 24 × 50 mm No. 1.5 coverslips (Fisherbrand cat. 12541033). Histological images of H&E-stained tissues were captured using an Olympus BX43F microscope with a 40× objective. Low-power 4× imaging was used only for whole-section overviews and region of interest (ROI) delineation; all quantitative analyses were performed exclusively using high-power 40× or 60× objectives with appropriate numerical apertures. For high-resolution imaging, z-stacks spanning the full 40-μm section thickness were acquired using 0.3-0.5 μm optical steps, ensuring Nyquist sampling. All quantification relied solely on these high-power, z-stacked datasets.

**Purkinje cell quantification** For Purkinje cell quantification, cerebellar sections from P11 and P45 mice were stained for Calbindin. Images were acquired on an Olympus BX43F microscope using a 40× objective (NA 0.95), and background and noise were reduced using CellSense background correction. PCs were manually counted in FIJI/ImageJ using the Cell Counter plugin in 3-4 representative midsagittal vermal regions per animal by scorers blinded to group identity. ROIs were predefined using consistent anatomical landmarks and maintained at identical field dimensions across all sections, animals, and experimental groups, ensuring that PC counts were obtained from equivalent tissue areas.

**Purkinje cell dendritic morphology** PC dendritic complexity and dendritic length were quantified using Neurolucida 360 (MBF Bioscience). Confocal images were acquired with an Olympus FV1000 using a 60× oil immersion objective (NA 1.4; x–y resolution: 0.124 μm/pixel; z-step: 2.0 μm). Dendrites were traced semi-automatically, and complexity was calculated as:[(terminal order sum + number of terminals) × total dendritic length / number of primary dendrites]. For each mouse, mean dendritic metrics were derived by averaging across 20 traced PCs, with $n$ = 4–5 animals per group.

**Granule cell quantification and granule layer area measurements** Granule cell quantification was performed on H&E-stained 40× images using the same blinded manual counting procedure within standardized ROIs described above. External granule layer (EGL) and internal granule layer (IGL) cross-sectional areas were quantified on H&E-stained 40× images using QuPath.

**P21 and P27 immunofluorescence quantification** P21 and P27 immunofluorescence quantification was performed on cerebellar sections from the relevant developmental time points. Images were acquired under consistent imaging conditions across groups, and P21- or P27-positive cells were manually counted within predefined ROIs by investigators blinded to experimental condition. ROIs were selected using consistent anatomical landmarks and maintained at identical field dimensions across all sections. Counts were averaged at the animal level and used for statistical analysis.

**VGLUT2 puncta quantification** VGLUT2 puncta quantification was conducted on four midsagittal sections per animal, using two standardized ROIs per section located in cerebellar lobule III and encompassing the molecular layer and PC layer, yielding eight images per animal. Images were acquired with a 60× oil immersion objective (NA 1.4). ROIs were manually delineated in QuPath, and puncta were quantified using a combined manual and semi-automated threshold-based approach performed entirely in a blinded manner. Total puncta counts were normalized to ROI area, and animal-level means were used as the biological unit for statistical analysis.

**Statistical unit, blinding, and consistency across analyses** Across all endpoints, Purkinje and granule cell counts, P21/P27-positive cell counts, dendritic measures, granule-layer area, and synaptic puncta, the individual animal served as the experimental unit. Group identity was coded by an investigator independent of data collection and decoded only after quantification was complete. Experimenters could not be blinded during hypoxia, which requires direct manipulation of the chamber; for all other procedures, samples, images, and recordings were processed blind to group. All histological, immunofluorescence, electron-microscopy, and behavioral scoring, including ROI selection and quantification, was performed blind, and automated or semi-automated pipelines (QuPath, Neurolucida 360, DeepLabCut/B-SOiD, Seahorse Wave) were applied identically across coded samples.

**Human postmortem cerebellar tissue collection and neuropathology** A retrospective neuropathological analysis was performed on human postmortem cerebellar tissue from eight subjects, including four preterm and four term infants. Samples were obtained from the pathology department at Children's National Hospital, and demographic and clinical details are summarized in Supplementary Tables 2-5. Postmortem subjects were selected from institutional archives based on gestational age, postmenstrual age at death, tissue availability, and preservation quality, with exclusion of major brain-destructive pathology and documentation of relevant clinical comorbidities. Cerebellar tissue was processed as formalin-fixed paraffin-embedded material using standardized histopathology procedures. Sections were cut at 5 μm thickness and stained with H&E. Cellular morphology and cytoarchitecture were assessed within the EGL, IGL, molecular layer, and PC layer. Quantification mirrored the murine neuropathology pipeline, using anatomically matched ROIs of equivalent size for layer cross-sectional area measurements and cell counts, with all analyses performed blinded to group identity.

#### Bulk RNA sequencing

Cerebellar tissue was collected for all four mouse groups (Sal-N, Sal-Hx, LPS-N, LPS-Hx) with $n$ = 3-5 for P11 mice. The tissue was then sonicated with the Ultrasonic Processor (Cole-Parmer, serial number: 2011040615) using QIAzol Lysis Reagent (QIAGEN, cat. 79306) and sent to Novogene Corporation Inc. (Durham, NC) for RNA sequencing. In brief, Qubit and Bioanalyzer instruments were used to perform a quality control check for RNA samples. The NEBNext Ultra II non-directional RNA Library Prep kit was used to prepare libraries for sequencing. Labchip and qPCR assays were used to assess library quality and concentration. The Novoseq6000 instrument was used to perform PE150 sequencing of the libraries. RNA Integrity Number (RIN) values of the samples ranged from 9.1 to 9.8. Preprocessing of the data was performed using Salmon version 0.8.2 for alignment to the gencode vm10 reference index, and TXImport version 1.2.0 for transcript level summarization.

Human postmortem cerebellar tissue was collected after formalin fixation and paraffin embedding from Children's National Hospital Department of Pathology in accordance with IRB and institutional guidelines (IRB #00011850, IRB #00015350). Supplementary Tables 2-5 provide demographic data for the subjects. RNA extraction for human cerebellar tissue ($n$ = 4/preterm, n = 4/term) was performed utilizing the QIAzol RNeasy FFPE Kit [QIAGEN, cat. 73504]. Procedure for extraction is retrievable in https://www.qiagen.com/us/products/discovery-and-translational-research/dna-rna-purification/rna-purification/total-rna/rneasy-ffpe-kit.

#### Cross-species enrichment of insult-specific transcriptional signatures

Cerebellar bulk RNA-seq counts from the four P11 experimental groups (Sal-N, Sal-Hx, LPS-N, LPS-Hx) were modeled with DESeq2 (v1.50.2) using a multi-group design (~ group). Samples flagged as transcriptomic outliers on PC1–PC2 of the top-500-variable-gene PCA (silhouette < −0.5, with cross-clusters explainable by a shared LPS or shared hypoxia exposure retained as expected biology) were excluded prior to fitting. Three pairwise contrasts against the Sal-N reference were extracted: LPS-N vs Sal-N (MIA signature), Sal-Hx vs Sal-N (hypoxia signature), and LPS-Hx vs Sal-N (sequential MIA+Hx signature). For each contrast, genes were ranked by the absolute value of the Wald statistic, and the top 200 ranked genes were retained as the candidate insult signature. Insult-unique signatures were defined as genes appearing in only one of the three top-200 lists; each insult-unique set was split by the sign of its Wald statistic in the source contrast, yielding six P11 insult-unique signatures (MIA-up/down, Hx-up/down, MIA+Hx-up/down). Mouse Ensembl gene identifiers were mapped to 1:1 human Ensembl orthologs (17,187 retained); genes without a 1:1 ortholog were excluded from cross-species analyses. Differential expression in the human postmortem cerebellar cohort (Hu_Original, preterm vs term, n=8) was modeled with DESeq2 to generate per-gene Wald statistics, yielding a ranking of 19,849 testable genes used as the cross-species reference. Each insult-unique signature was tested for concordant enrichment in this ranking using competitive enrichment with CAMERA pre-ranked (`limma::cameraPR`, inter-gene correlation 0.01), with Benjamini–Hochberg correction applied across the six signatures; rank-shift effect size was quantified by Wilcoxon area under the curve (AUC) of set-gene positions in the human ranking, with 95% confidence intervals estimated from 1,000 bootstrap resamples. For the MIA+Hx insult-unique signature, a barcode plot displayed the position of set genes along the human Wald ranking (up-set in red above the axis, down-set in blue below), with kernel-density curves visualizing local enrichment and vertical reference lines at the 25% and 75% quantiles. All analyses were performed in R 4.5.3 (DESeq2 v1.50.2, limma v3.68.2, AnnotationDbi v1.72.0).

#### Spatial transcriptomics

A single Visium slide was used for mouse tissue, with two cerebellar tissue sections from each of four treatment groups on each capture area. Tissue was embedded in OCT, cryo-sectioned at 10 μm, and mounted onto Visium capture slides according to the 10x Genomics Visium protocol (retrievable in www.10xgenomics.com). The library construction was carried out according to the manufacturer's protocols. The captured tissue sections were lysed, and the RNA was extracted and reverse-transcribed to cDNA using the 10x Genomics Visium Spatial Gene Expression Reagent Kit. The cDNA was then amplified, and sequencing libraries were prepared. The libraries were then sequenced at Paired-End 150 bp on the Illumina NovaSeq6000, targeting 50,000 reads/spots. The raw data obtained from the sequencing were processed using the SpaceRanger (10x Genomics) to align and quantify gene expressions. The resulting expression matrix was imported into Seurat (version 4.0.1) for downstream analysis in R. The data were normalized using the NormalizeData function in Seurat with the LogNormalize method to account for differences in library size and to minimize the impact of technical noise. Dimensionality reduction was performed using the RunPCA function, followed by clustering using the FindClusters function. The results were visualized using ggplot2 (version 3.3.5) plotting functions. Pathology annotations were included in the cloupe file generated by the SpaceRanger pipeline. Clustering was performed to resolve the pathology annotations in accordance with gene expression using the FindClusters function in Seurat. The resulting clusters were further annotated based on gene expression profiles and pathology information, allowing for the identification of spatially distinct cell types and the characterization of their gene expression patterns in different pathologies. Differential expression analysis was performed using the pseudobulk approach with the FindMarkers function in Seurat, followed by batch correction with the RemoveBatchEffect function in Limma (version 3.48) and further analysis with the DESeq2 (version 1.34.1) package in R. Cell-type annotations were assigned by integrating pathology-guided spatial localization on H&E-stained sections with marker gene expression profiles identified through Seurat clustering, using cell-type-defining gene panels derived from the Aldinger et al. cerebellar single-cell atlas[27] as biological priors. Analyses were conducted in R version 4.1.

**CoGAPS patterns** We ran Coordinated Gene Activity in Pattern Sets (CoGAPS), as previously described in[28,29], on the aggregated set of mice Visium samples to identify cell-type- and spatially-organized transcriptional patterns. Spots were restricted to those annotated as cerebellar cortex folia, including the molecular layer, Purkinje cell layer, and internal granular layer. Regions identified as white matter were systematically excluded based on both histological assessment of tissue architecture and cell-type annotation profiles, thereby ensuring that all retained spots for downstream analysis corresponded specifically to the cortical laminar compartments of the cerebellum. Genes with zero expression across all remaining spots were removed, and remaining expression values were log1p-transformed prior to analysis.  CoGAPS (v3.24.0) was run in R 4.3.0 using 15 patterns, 20000 iterations, and seed = 123 on these log-transformed expression values. We used the genome-wide distributed mode with nSets = 5, which partitions the features into five subsets and performs a two-pass procedure internally. In the first pass, matrix decomposition was performed independently on the 5 feature subsets using independent Markov chains. Patterns identified from each subset were matched across subsets by correlation in sample space, and clustered to form consensus patterns.  In the second pass, the consensus pattern matrix was fixed, and the feature loadings were learned for each subset and stitched across subsets to obtain the final solution. Pattern stability was evaluated using the number of contributing sets and the correlation of each subset-derived pattern to the mean consensus pattern. Patterns that failed to meet the consensus-matching criteria were not used for downstream interpretation. In our analysis, 15 patterns were specified at initialization, and 13 robust patterns (M-1 through M-13) were retained in the final consensus solution.

**Gene set enrichment analysis (GSEA)** GSEA was performed independently for each CoGAPS pattern using the feature loadings as the ranking statistic. Enrichment was computed using fgsea (v1.28.0) in R (v4.3.0), with positive scoring (scoreType='pos'). Gene sets were obtained using the msigdbr package (v 24.1.0), selecting the C5 collection (Gene Ontology gene sets) for Mus musculus, excluding Human Phenotype Ontology (HPO) gene sets. Pathways were filtered based on an adjusted p-value cutoff of $< 0.05$ and gene set size between 15 and 500 genes. The top 25 pathways per pattern ranked by p-value were retained for visualization.

**LIANA cell-cell communication analysis** Spatial ligand–receptor communication between granule cells (GC) and Purkinje cells (PC) was inferred using the LIANA framework[30], applied to the Visium spatial assay after Seurat-based annotation as described above. Spots were restricted to the PC and GC compartments, and genes with no detected counts were removed. LIANA's Consensus ligand–receptor resource was converted to mouse orthologs (NCBI taxon 10090) and used for all analyses. To preserve biological replication, LIANA was run independently on each sample (two biological replicates per condition); samples lacking either cell type or with fewer than five cells of either type were excluded. Within each sample, data were log-normalized (NormalizeData) and ligand–receptor interactions between PC and GC were scored with liana_wrap (minimum of five cells per identity) and combined across LIANA's individual methods into a consensus rank using liana_aggregate. Per-sample results were pooled, and interactions were retained only if present in both replicates of a condition; their interaction magnitude (sca.LRscore) and specificity (NATMI edge specificity) were averaged across replicates to yield condition-level estimates. For visualization, interactions were prioritized by their mean aggregate rank across replicates, and the top 10 in each direction (PC-to-GC and GC-to-PC) were displayed on shared magnitude and specificity scales (Supplementary Fig. 5A–D).

**Tricycle** We applied Tricycle analysis as previously described in[31] to evaluate cell-cycle phases at the spot level using Visium spatial transcriptomic data. Raw counts were log1p-transformed, and Tricycle (v 1.10.0) was applied in two complementary ways. First, a continuous cell-cycle position, was estimated by projecting the spot-level profiles into the pre-established Tricycle reference embedding. This projection maps each cell onto the cyclical trajectory based on the similarity of its gene expression profile to the reference, assigning a cell cycle position as an angle ranging from 0 to $2\pi$. Separately, discrete cell-cycle stages (G1/S, S, G2, G2/M, M/G1) were assigned to each spot using the Schwabe stage estimator as reimplemented in Tricycle, which assigns each spot based on the expression levels of phase-specific marker genes, ensuring accurate and biologically meaningful categorization[32].

#### Electron microscopy

For ultrastructural assessment of the alterations in Purkinje and granule cells in the cerebellar cortex, mice (P11 and P45) from all experimental groups (Sal-N, Sal-Hx, LPS-N, and LPS-Hx) were transcardially perfused with 4% PFA and 0.5% glutaraldehyde in 0.1 M PBS. After overnight postfixation, vibratome-sagittal sections of the cerebellum were cut at 300 µm and incubated in 3% potassium ferrocyanide dissolved in 0.3 M cacodylate buffer containing 4 mM calcium chloride and 4% aqueous osmium tetroxide (for 1 hour at 4°C). This was followed by a filtered thiocarbohydrazide solution incubation for 20 minutes at room temperature. Then the sections were washed and incubated in 2% aqueous osmium tetroxide solution (30 minutes at room temperature). The sections were placed overnight in 1% uranyl acetate at 4°C and stained en bloc with Walton's lead aspartate solution. The samples were dehydrated using a graded ethanol series (50%, 70%, 85%, 95%, 100%) and then placed in propylene oxide before being infiltrated with EPON resin (EMbed-812; Electron Microscopy Science, Hatfield, PA, USA). The samples were embedded between thermoplastic fluoropolymer films (Aclar, EMS) and polymerized in an oven at 60°C for 48 hours. Blocks containing the cerebellar lobules were trimmed/mounted and sectioned with an ultramicrotome at 120 nm (UC7 Leica Microsystems). The ultrathin sections were placed on silicon wafers and carbon-taped onto aluminum stubs for Scanning Electron Microscopy imaging using a Helios NanoLab 660 FIBSEM (ThermoFisher). A concentric backscattered electron detector (CBS) in immersion mode was utilized at a 4 mm working distance, using 2 kV and 0.40 nA electron probe to increase the backscattered electrons density to the detector. Low magnification (1000x) overview images of the entire cerebellum were taken to target specific regions of interest for high-resolution imaging (cell layers). Subsequently, high-resolution tile image sets of the specific areas containing randomized cell types (Purkinje and granule) were acquired and stitched at 80,000x magnification (5s dwell time, 3072x2048 resolution) with a pixel size of 1.6862 nm (MAPS 3.22, ThermoFisher).

**2D mitochondrial segmentation and morphological assessment in Purkinje and granule cells** To quantify differences in cerebellar mitochondrial ultrastructure between all experimental approaches, we converted 2D tile-Scanning Electron Microscopy images to a native.sis file in arivis Vision4D software (Zeiss). Then, using the analysis panel (arivis), we built a pipeline for mitochondrial segmentation analysis. In summary, we segmented (manually) all mitochondria in each cell (region of interest), tagging them as individual objects. Quantitative object data for each segmented mitochondria in the ROI, including morphological properties such as 2D area, surface area, short side, long side, and density (number of mitochondria per $mm^2$), were obtained through arivis operation and exported to Excel. Statistical analysis was performed using variance analysis followed by post-hoc analysis with Dunn’s multiple comparisons tests. The confidence of $P<0.05$ was considered statistically significant. Statistical analyses were performed at the biological scale appropriate to each measurement. Mitochondrial surface area was analyzed at the level of the individual mitochondrion, capturing the distribution of organelle sizes within each cell population[33], whereas mitochondrial density was analyzed at the level of the cell. Per-animal summary statistics for all mitochondrial measures are provided in the supplementary material. For mitochondria health analysis, two blinded assessors independently performed mitochondrial scoring measurements. Mitochondrial cristae appearance, matrix, and vacuolization were graded by using a five-point scale: 5 for the most degenerated or absent appearing cristae with mitophagy appearance and 1 for the most intact and well-ordered cristae. Observers were trained to use the scale and checked each other over one negative per animal to ensure conformity. See Fig. 4 for the scale. The unit of statistical analysis was matched to the biological scale of each measurement. The exact number of cells and mitochondria analyzed for each group is provided in Supplementary Table 6.

#### Seahorse XF assessment of mitochondrial function

We isolated cerebellar tissue from mice from all four groups (Sal-N, Sal-Hx, LPS-N, LPS-Hx). Tissue was used immediately to preserve the native metabolic microenvironment of the cerebellum, providing a systemic readout of group-level metabolic state. Tissue sections of 100 µm depth and 2 mm diameter were obtained and placed in artificial cerebrospinal fluid. Cerebellar tissue was then added to an islet capture microplate to measure mitochondrial respiration in the Seahorse XFe24 Analyzer© (Agilent Inc., Santa Clara, CA). The Seahorse XF Analyzer© utilizes inert optical microsensors to measure oxygen consumption rate (OCR). OCR indicates mitochondrial oxygen consumption by oxidative phosphorylation to generate adenosine triphosphate (ATP). Mitochondrial respiration was assessed using a standard Mito Stress Test protocol[34] consisting of sequential injections of oligomycin (1 µM), FCCP (3 µM), and a combined rotenone/antimycin A solution (5 µM total), each delivered in 25 µL injection volumes. Proton leak was calculated as oligomycin-inhibited OCR minus non-mitochondrial respiration (antimycin A/rotenone-inhibited OCR), and coupling efficiency was derived as the proportion of basal respiration used for ATP synthesis. The sample size for each group is as follows: P11 Sal-N ($n$ = 8), P11 Sal-Hx ($n$ = 14), P11 LPS-N (*n* = 8), P11 LPS-Hx (*n* = 10); and P45 Sal-N (*n* = 6), P45 Sal-Hx (*n* = 5), P45 LPS-N (*n* = 6), P45 LPS-Hx (*n* = 6).

#### Mouse Behavior

Behavioral testing was conducted during the light phase of the circadian cycle, between the hours of 8 AM and 12 PM, in facilities maintained by the Neurobehavioral Evaluation Core of the Intellectual and Developmental Disabilities Research Center at Children's National Hospital. Mouse behavioral assays were conducted by the research team after receiving training by the Core staff. On the days of testing, mouse cages were habituated to the testing environment: cages were brought into the testing room 20-30 minutes before the start of each behavioral assay.

Per-group sample sizes for behavioral analyses are reported after application of *a priori* exclusion criteria specific to each assay and analysis pipeline. All mice underwent the same behavioral testing battery, including locomotor assessment on the Erasmus Ladder, the Social Interaction Test (SIT), and Open Field Testing (OFT). Behavioral datasets were curated to preserve matched cohorts across assays; however, exclusions were required in select cases because of incomplete recordings, faulty video acquisition, missed tracking intervals, or insufficient tracking quality. For tracking analysis (OFT and B-SOiD) specifically, additional exclusion criteria were applied related to frame completeness, tracking continuity, and model-training requirements, as variability in frame fraction and pose-estimation quality can substantially affect unsupervised behavioral classification performance.

Final sample sizes following assay-specific quality control were as follows: Erasmus Ladder locomotor analysis, *n* = 28 mice; SIT, *n* = 44 mice; OFT analyses, *n* = 24 mice; and B-SOiD behavioral classification, *n* = 44 mice. Group-specific sample sizes for each assay are provided in Supplementary Table 7.

**Locomotor function and adaptive learning** Locomotor testing was assessed with an ErasmusLadder apparatus (ERLA-0010; Noldus Information Technology bv Wageningen, The Netherlands). ErasmusLadder is a fully automated behavioral assessment tool with specificity for cerebellar functions[35,36,37]. The equipment is comprised of two goal boxes linked by a horizontal ladder with pressure-sensitive rungs. Each of the rungs is connected to a central integrated processor. When a mouse steps on a rung, the pressure on the rung activates a sensor that logs each step into the processor. As the mouse walks along the entire horizontal ladder, the rungs on which the mouse has stepped are registered into the processor, and the stepping patterns of each mouse are generated. The ErasmusLadder system can generate a wide range of metrics to analyze locomotor coordination and adaptive cerebellar learning. In particular, the ErasmusLadder software can be used to measure the following metrics:

1. Missteps: the percentage of steps whereby a mouse makes a faulty movement onto the lower misstep rungs, rather than the default walking rungs.
2. Backsteps: the percentage of steps whereby a mouse makes a backward movement, i.e., in the direction opposite to that of the tailwind.
3. Short steps and long steps: the percentage of steps whereby a mouse makes a movement from one rung to the next rung on the same side (short step) or the second next rung (long step).
4. Pre-perturbation step-time: the time-difference between subsequent rung activations immediately preceding the obstacle on the same side. The obstacle consists of an additional rung with an automated activation system that blocks the normal path of the mouse in the horizontal ladder.

5. Post-perturbation step-time: the time-difference between rung activation immediately preceding the obstacle and immediately following the obstacle on the same side. Both pre- and post-perturbation step times are measured in milliseconds.

The Erasmus Ladder software (versions 1.0 and 1.1) was used to operate the ladder system under its default protocol for assessing cerebellar function at postnatal days 35-42 ± 2 (P35-42). The experimental design for these assessments involved the standard protocol of 4 training sessions followed by 4 challenge sessions conducted over consecutive days. Training sessions included 42 unperturbed trials per session, during which no obstacles were presented. In contrast, challenge sessions consisted of 42 trials presented in a randomized sequence across three categories: unconditioned stimulus-only (US-only) or obstacle, conditioned stimulus-only (CS-only) or auditory cue, and paired trials. During US-only trials, the animals encountered a computer-controlled obstacle that unpredictably obstructed their movement along the ladder. A high-pitched warning tone was delivered at random points along the ladder in CS-only trials. For paired trials, the conditioned stimulus was followed by the unconditioned stimulus with a fixed interstimulus interval of 250 milliseconds. "Post-perturbation step-time" for sessions 1 through 4 (where there is no physical object perturbation) was measured based on the first step, which involves computer selection of which obstacle rung along the ladder would be activated but without activating and presenting it. This metric serves as an "internal control" to compare post-perturbation step-time measurements across all 8 sessions. For locomotor behavioral experiments, outliers were excluded from group analyses based on two criteria: (1) mice with >25% jumps on any of the last four sessions were excluded from analysis; (2) for post-perturbation step-time, Robust regression and Outlier removal ROUT method (Q=1%) was used to screen outliers.

**Social Interaction Testing and Open Field Testing** Sociability was evaluated with the three-chambered social approach test[38,39]. A modified three-chamber apparatus was used. During the initial habituation phase, the subject mouse was first confined to the middle chamber for 10 minutes, followed by 10 minutes of free exploration across all three empty chambers. This second habituation period, in which the mouse was already acclimated to the apparatus and no social or object stimuli were present, was used to assess open-field behavior. Testing and scoring were performed blind to group, and the apparatus was cleaned with 10% ethanol between subjects. The test session then consisted of a 10-minute period in which a novel cylinder object was placed in one side chamber, while a novel mouse was placed inside a similar cylinder in the opposite side chamber, allowing the subject mouse to freely explore all chambers. The novel mouse was carefully selected to be older and calmer than the test subjects and was kept consistent for all animals. A single adult female mouse was used as the novel stimulus across all test animals to ensure consistent, non-aggressive social stimulation regardless of subject sex, as female stimulus mice do not provoke territorial or agonistic responses from either male or female subjects[40]. Mouse movements were captured on a FLIR Camera with Computer CS-Mount 1.8-3.6mm Varifocal lens. Behavioral/movement tracking was performed through a customized Bonsai workflow[41]. Subject mice were at age P44 ± 2 days at the time of testing. Group sizes were $n$ = 6-8 per treatment group.

**DeepLabCut (DLC) pose estimation** We used DeepLabCut v2.1[42,43] to detect and quantify movement data in the three-chamber social behavior test and the prior habituation phase. We labeled 12 body parts of the subject mouse, four points on the stranger mouse cage (as a reference), and four points on the novel object in a total of 100 images from five videos (20 images/video; one video/animal; four videos of the test-phase and one video of the acclimation phase) across different experimental groups to generate input data for training. The ResNet-50-based neural network was used for training. Default configurations were retained in the pipeline, including a training fraction of 0.95. We trained the model for 30000 iterations. The test-error with a p-cutoff of 0.6 was 4.07 pixels. We used this model to predict pose estimation on all social behavior videos.

**B-SOiD behavior segmentation** Prematurity is linked to subtle, long-term motor deficits that may escape detection by standard behavioral assays[44]. Detailed kinematic analysis offers a sensitive means to uncover these impairments by quantifying fine locomotor features[45,46,47]. In mouse models, such analysis can reveal lasting neuromotor disruptions stemming from early-life insults that affect cerebellar or cortical development. We used B-SOiD[48] to identify detailed movement patterns during the sociability test, with the goal of uncovering subtle changes in behavior linked to differences in social interaction. Segmentation of pose estimation datasets of social behavior was generated from DeepLabCut. We used the streamlit app to run B-SOiD as detailed on the B-SOiD Github repository (https://github.com/YttriLab/B-SOID). For B-SOiD training, we used a dataset of ten videos analyzed by DeepLabCut. 0.5%-1.5% of video length was used for B-SOiD cluster identification. B-SOiD identified 14 kinematic patterns, that were distinctly clustered and identified using this configuration in the HDBSCAN assignment. Supplementary Table 8 provides a description of the kinematic patterns identified. All 14 clusters identified by B-SOiD were retained for analysis. Detailed reporting focuses on clusters demonstrating statistically significant between-group divergence. Random Forest classifier cross-validation accuracy was ~0.89 based on 20% of the data. We used this B-SOiD model to segment behavior across all DeepLabCut analyzed videos of social behavior.

**DLC and B-SOiD analysis and statistics** We wrote a custom python script to divide the chamber in six grids, which allowed us to identify direct time spent in contact with the novel mouse, direct time spent in contact with the novel object, and time spent exploring in the remaining grids. We used each chamber border and a midline point to create the grids. The custom Python script allowed us to analyze first-order data including time spent in each specific grid from social behavior videos. Between-group comparisons on first-order data were conducted using pairwise two-sample Welch's t-tests, while within-group comparisons (social versus object chamber) were performed using paired Wilcoxon signed-rank tests. The same code was used to build assessing areas in the chamber corners and chamber gates, and to analyze the time spent in corners versus open field during the 10-minute open field assessment. We wrote custom Python scripts to analyze B-SOiD segmentation groups and perform statistics. Since some videos recorded for social behavior had different frame rates than those used for B-SOiD model training, we excluded these videos from B-SOiD analysis. We used Wasserstein distance from scipy.stats and the statsmodels.stats.multitest library in a custom Python script to compare the Wasserstein distance between empirical cumulative distribution functions (CDFs) of successive B-SOiD groups across experimental groups with permutation tests. Benjamini–Hochberg correction was applied to *P*-values. For the heatmap comparing B-SOiD group distributions, we generated linkage matrices using linkage from the scipy.cluster.hierarchy library. Scripts are included in the Supplementary material.

#### Statistical analysis

Data were compiled in Microsoft Excel 16.85, and statistical analyses were performed and figures generated in R version 4.5.3 (packages: rstatix, car, emmeans, FSA) and Python 3.12.12 (scipy, statsmodels; matplotlib/seaborn). All graphs display mean ± SEM; all tests were two-tailed with significance set at $\alpha = 0.05$. Data were assessed for normality using the

Shapiro–Wilk test and for homogeneity of variance using Levene's test. When both assumptions were met (Shapiro–Wilk $P > 0.05$; Levene's $P > 0.05$), parametric analyses were performed using one-way ANOVA with appropriate post hoc comparisons, including Tukey's correction for multiple comparisons or uncorrected Fisher's least significant difference (LSD) for pre-planned pairwise contrasts (identified in the corresponding figure legends). When normality was violated across groups, non-parametric testing was conducted using the Kruskal–Wallis test with Dunn's multiple comparisons (Bonferroni-adjusted). For human cerebellar comparisons (term versus preterm), two-tailed Mann–Whitney *U* tests were used. In cases where normality was violated in one or two groups, but variances remained homogeneous, ANOVA was retained given its established robustness to moderate deviations from normality under equal variances. For motor behavioural data, comparisons were made between experimental groups using two-way repeated-measures ANOVA with Tukey's multiple comparisons test for post-perturbation step-times and missteps; the full ANOVA output is provided in Supplementary Table 9. Open-field, social-chamber, kinematic (B-SOiD) and cross-species transcriptomic comparisons used the tests specified in the corresponding *Methods* subsections (two-sample Welch's t test; paired Wilcoxon signed-rank test; Wasserstein-distance permutation testing with Benjamini–Hochberg correction; and CAMERA pre-ranked enrichment, respectively). Exact *P* values are reported in the figure legends. Sample sizes for histological, immunofluorescence and ultrastructural cohorts were estimated using the resource-equation approach for exploratory animal experiments, which targets error degrees of freedom in the range of ~10–20; for a four-group design this corresponds to a minimum of ~4 animals per group, consistent with the 4–6 animals per condition used here, while behavioural cohort sizes are reported per assay in the Methods and figure legends. The degree of statistical significance was denoted using asterisks: $^*P < 0.05$; $^{**}P < 0.01$; $^{***}P < 0.001$; $^{****}P < 0.0001$.

Supplementary Fig. 1

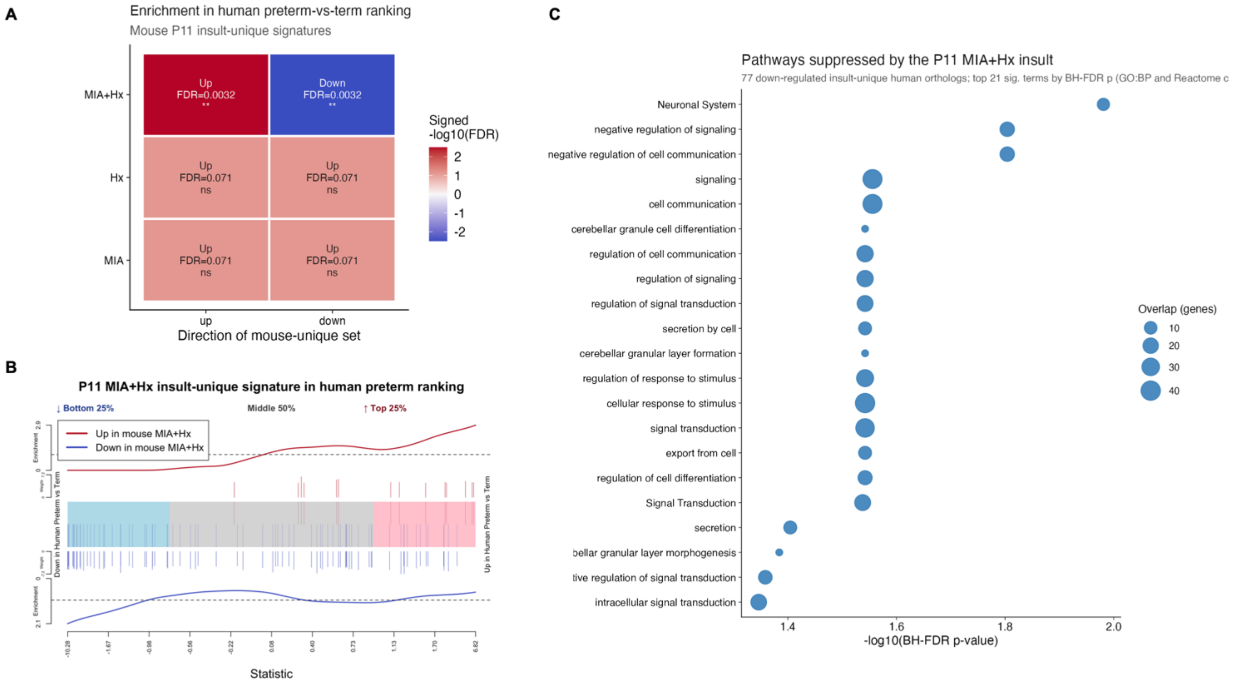

Supplementary Fig. 2

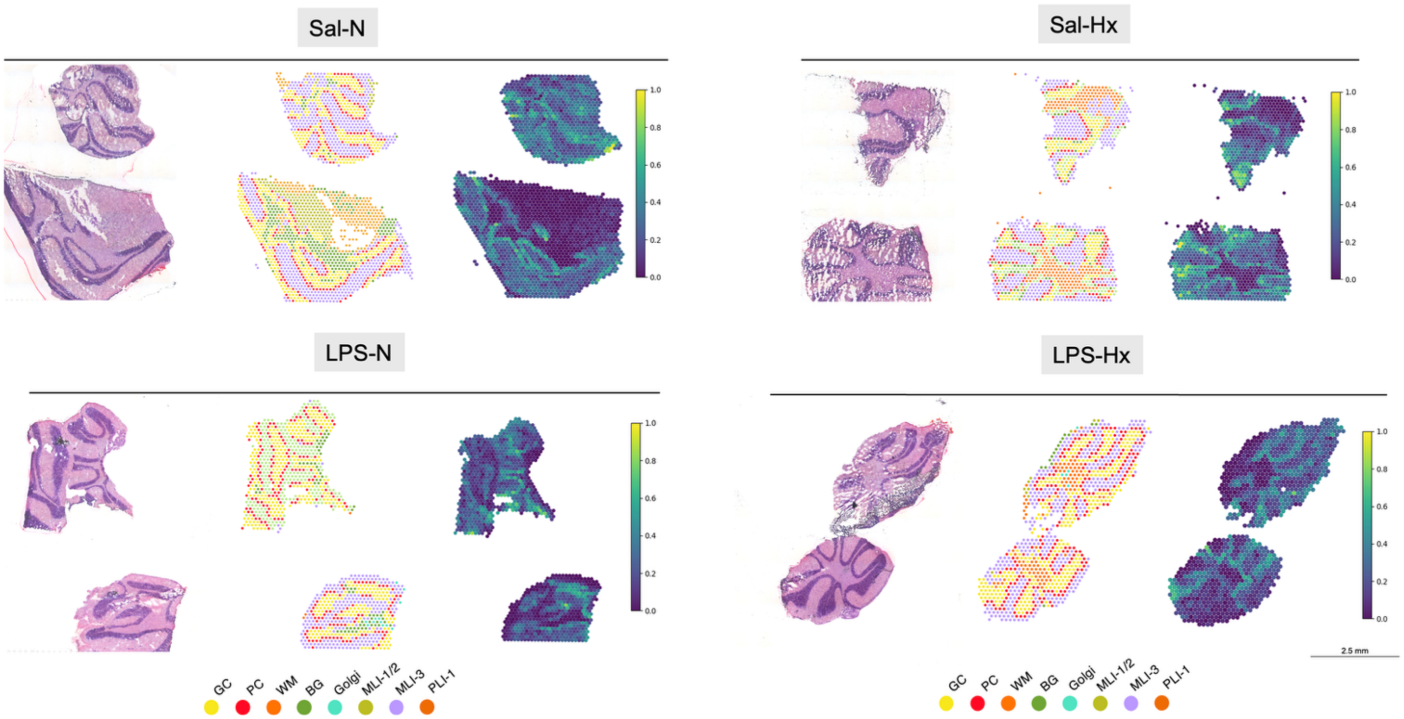

#### Supplementary Fig. 3

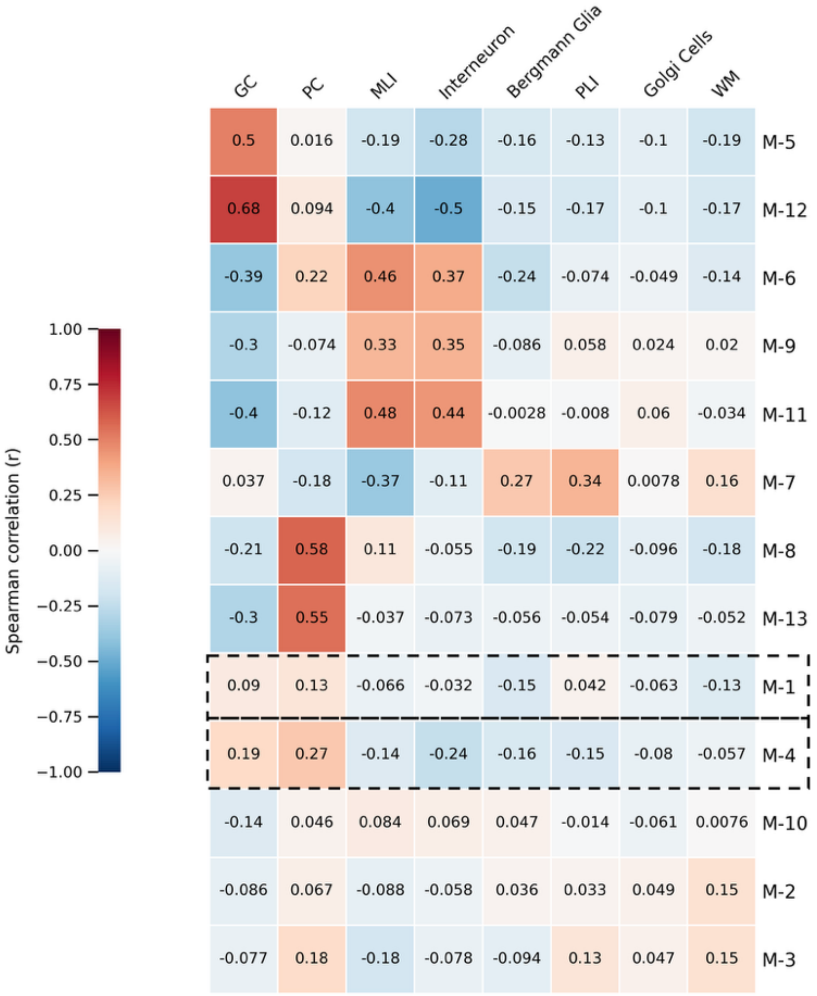

Supplementary Fig. 4

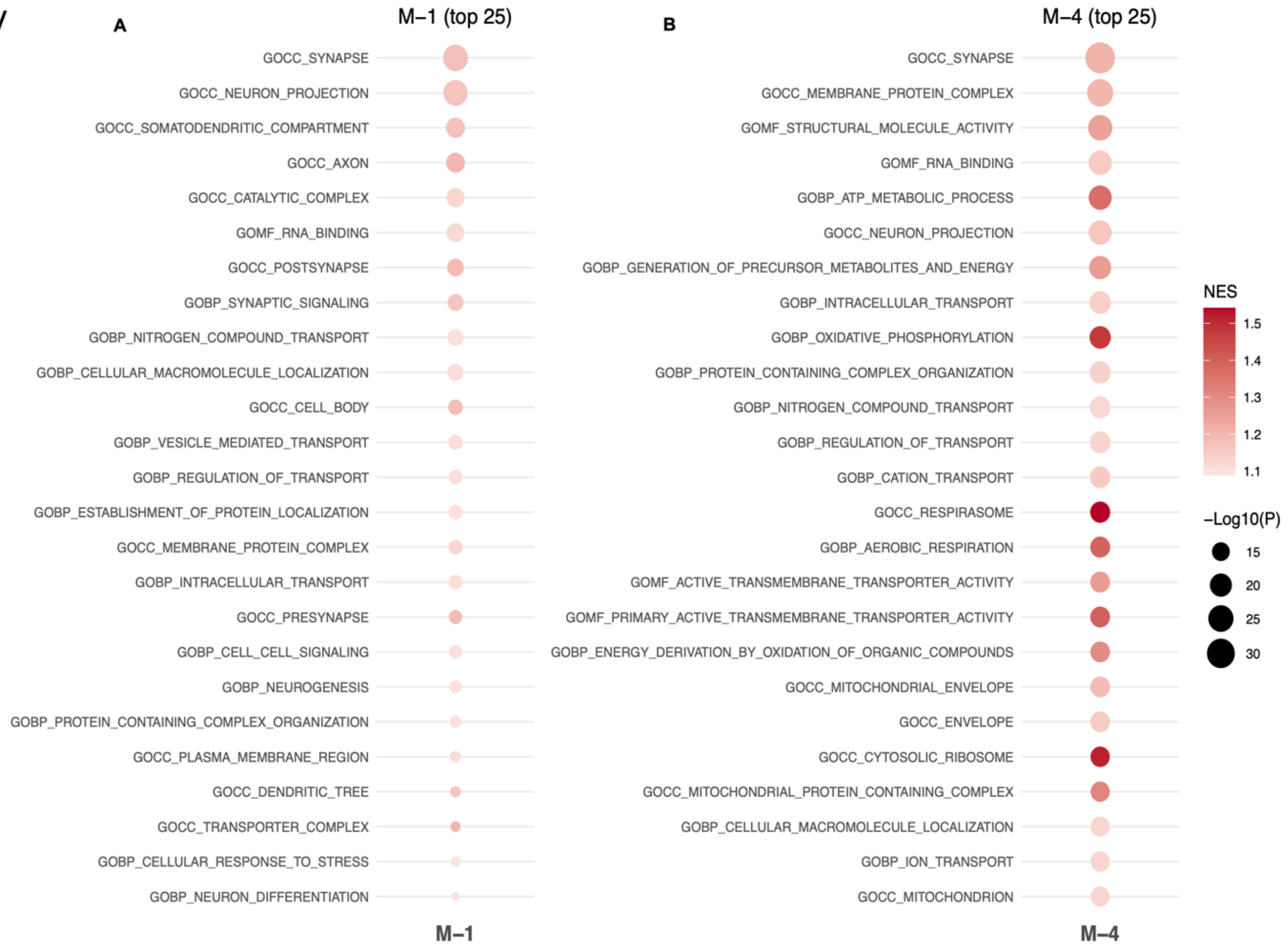

### Supplementary Fig. 5

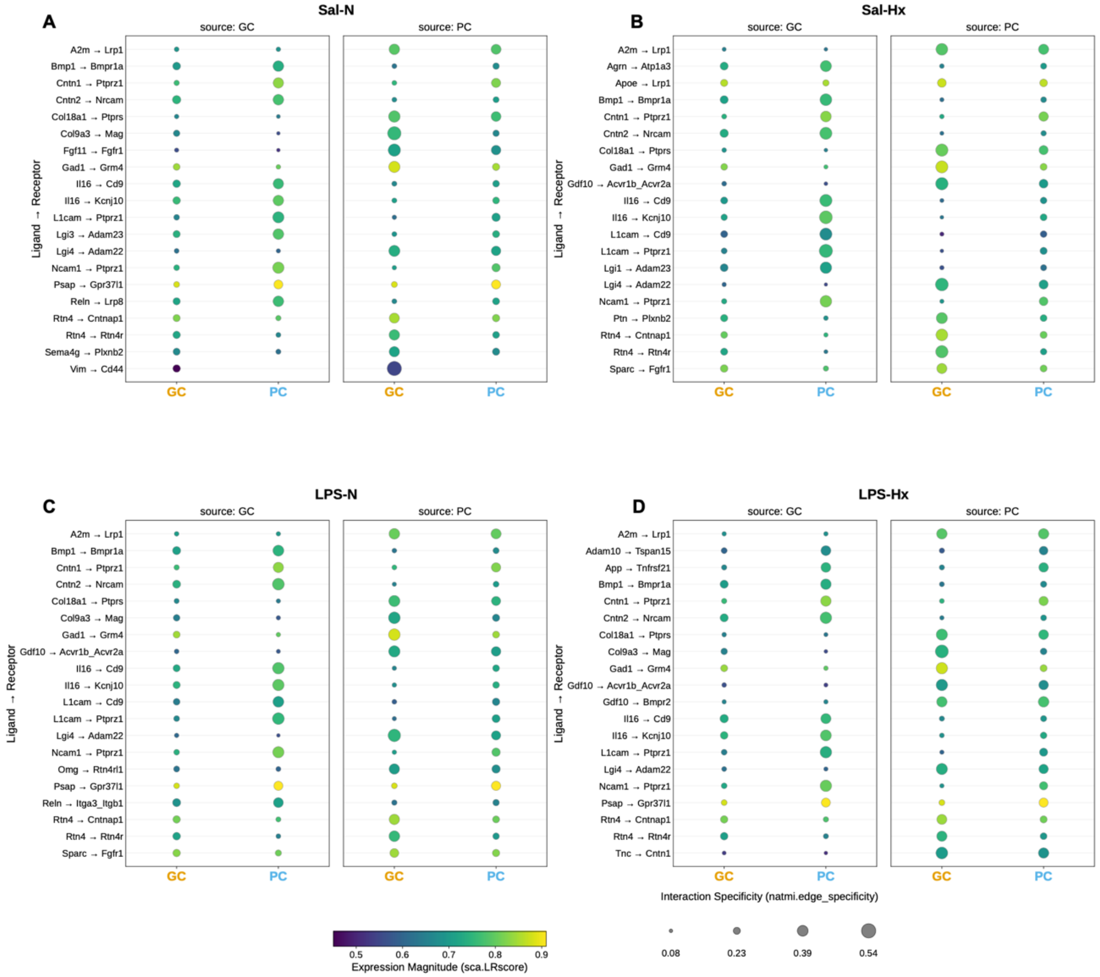

Supplementary Fig. 6

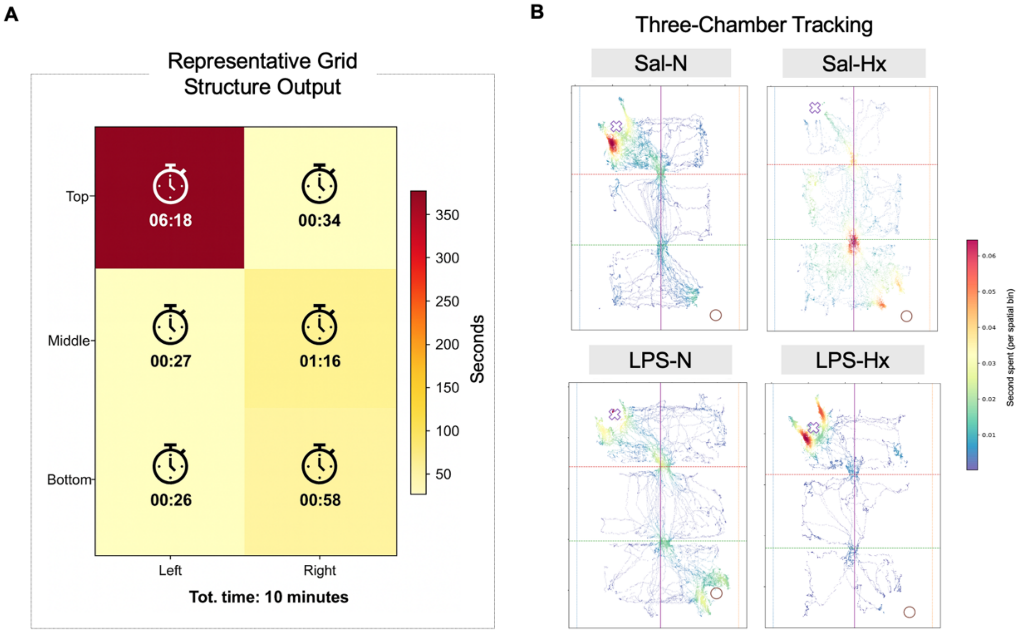

Supplementary Fig. 7

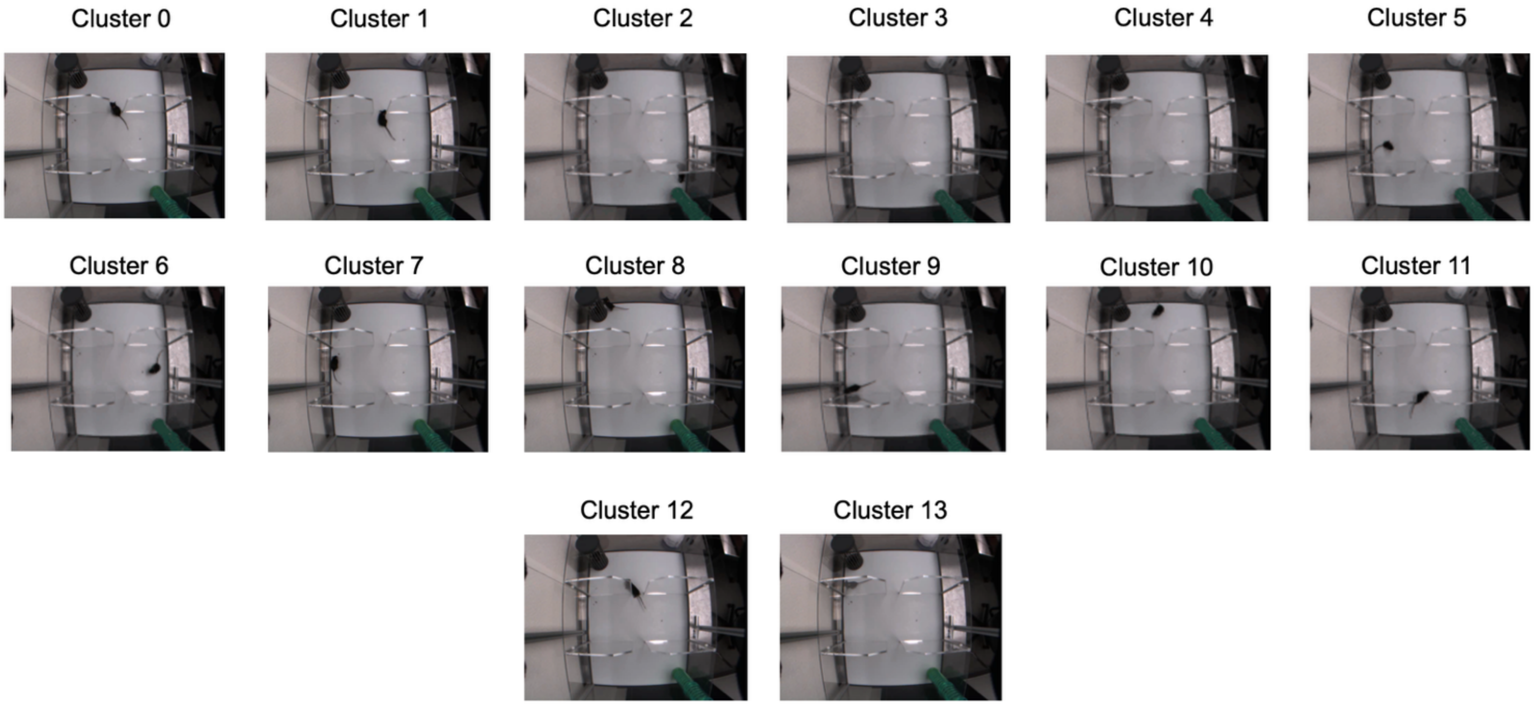

Supplementary Fig. 8

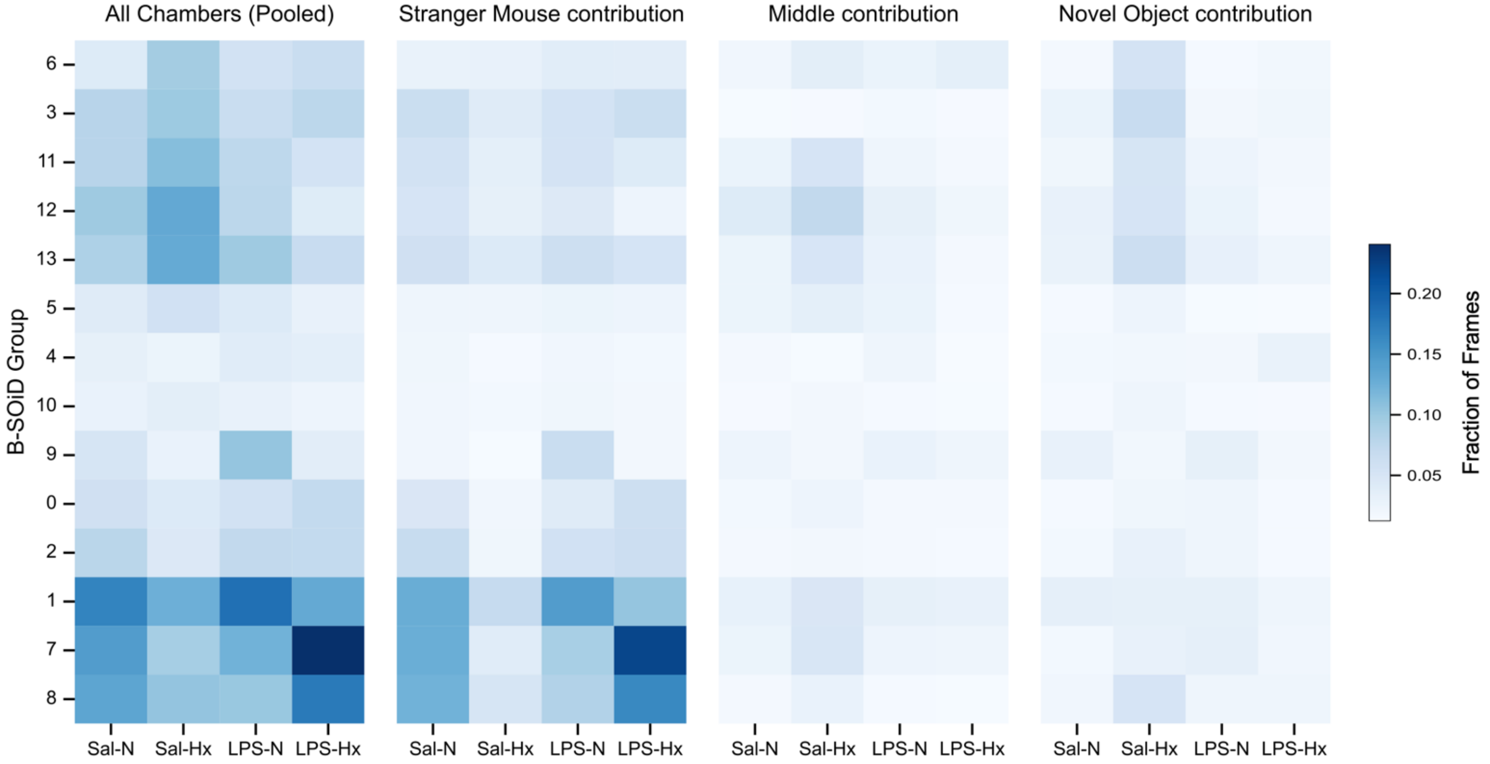

Supplementary Fig. 9

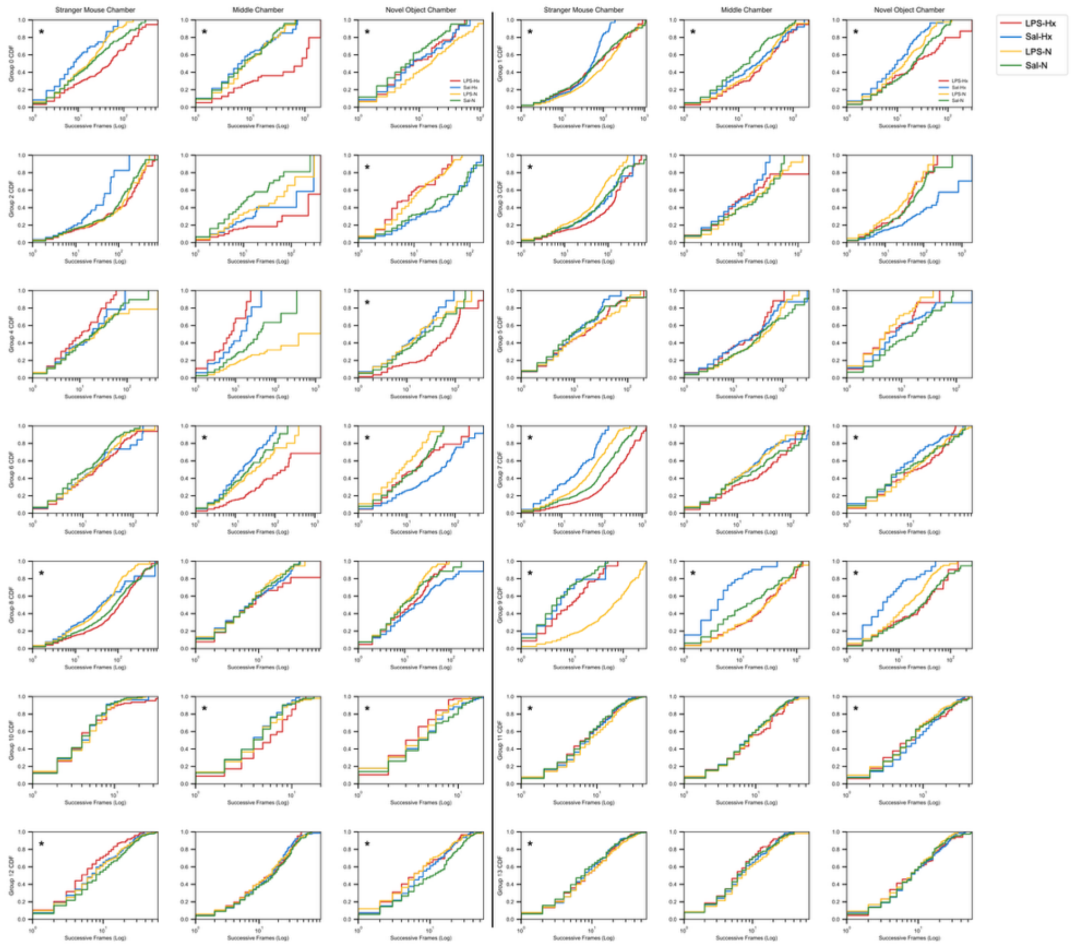

**Supplementary Fig. 1. Insult-specific cross-species enrichment of mouse P11 cerebellar insult signatures in human preterm cerebellum. (A)** CAMERA pre-ranked enrichment of mouse P11 insult-unique gene sets (rows: LPS-N vs Sal-N, MIA; Sal-Hx vs Sal-N, Hx; LPS-Hx vs Sal-N, MIA+Hx), split by direction of the mouse-unique set (columns), tested against the human preterm-vs-term Wald-statistic ranking. Tile colour: signed $-\log_{10}$(BH-adjusted CAMERA *p*); tile text: CAMERA direction, FDR, and significance (* < 0.05; ** < 0.01; *** < 0.001; ns ≥ 0.05). **(B)** Barcode plot for the MIA+Hx insult-unique signature on the human ranking (sorted left-to-right from most down- to most up-regulated in preterm). Background bands mark bottom-quartile (blue), middle 50% (grey), and top-quartile (pink) of the human ranking. Red ticks/density worm: mouse MIA+Hx-up set ($n$ = 14 orthologs; FDR = $3.2 \times 10^{-3}$, AUC = 0.77). Blue ticks/density worm: MIA+Hx-down set ($n$ = 77; FDR = $3.2 \times 10^{-3}$,

AUC = 0.42). Dashed lines: 25th/75th percentiles. Concordance of both directions indicates that the sequential signature recapitulates the human preterm transcriptome; MIA-only and Hx-only signatures are non-significant (FDR = 0.071). **(C)** Dot plot displaying pathways suppressed by the P11 MIA+Hx insult, derived from the 77 down-regulated insult-unique human orthologs. The top 21 significant terms by BH-FDR p-value are shown, drawn from Gene Ontology Biological Process (GO:BP) and Reactome databases. The x-axis represents $-\log_{10}$(BH-FDR *p*-value) and circle size represents the number of overlapping genes, ranging from 10 to 40. Enriched terms include pathways related to signaling, cell communication, signal transduction, cerebellar granule cell differentiation, cerebellar granular layer formation, regulation of response to stimulus, and neuronal system. For analysis, *n* = 3-5 per mouse group, and *n* = 4 human preterm and 4 human term subjects. Abbreviations: AUC, area under the curve; BH, Benjamini-Hochberg; CAMERA, Correlation Adjusted MEan RAnk gene set test; FDR, false discovery rate; GO:BP, Gene Ontology Biological Process; Hx, hypoxia; LPS, lipopolysaccharide; MIA, maternal immune activation; N, normoxia; ns, not significant; Sal, saline.

**Supplementary Fig. 2. Spatial Annotation of Cerebellar Cell-Type Domains Across Experimental Groups.**

Hematoxylin and eosin-stained mouse cerebellar whole sections, corresponding spatial cell-type annotation maps, and representative CoGAPS expression pattern maps are shown for Sal-N, Sal-Hx, LPS-N, and LPS-Hx samples. For each group, two biological replicates are displayed. Left column: hematoxylin and eosin-stained histological sections. Middle column: spatial spots colored by annotated cerebellar cell-type identity, including granule cells (GC), Purkinje cells (PC), white matter (WM), Bergmann glia (BG), Golgi cells, molecular layer interneuron subtypes (MLI-1/2 and MLI-3), and Purkinje layer interneurons (PLI-1). Right column: representative CoGAPS expression pattern maps, with expression intensity displayed on a continuous scale ranging from 0.0 (dark purple) to 1.0 (yellow). The paired histological, cell-type annotation, and CoGAPS pattern maps illustrate the anatomical distribution of major cerebellar cell populations and spatially resolved gene expression programs used for downstream spatial transcriptomic analyses. For spatial analysis, *n* = 2 per group. Scale bar, 2.5 mm. Abbreviations: BG, Bergmann glia; CoGAPS, Coordinated Gene Activity in Pattern Sets; GC, granule cell; Hx, hypoxia; LPS, lipopolysaccharide; MLI, molecular layer interneuron; N, normoxia; PC, Purkinje cell; PLI, Purkinje layer interneuron; Sal, saline; WM, white matter.

**Supplementary Fig. 3. Cell-Type-Specific Enrichment of CoGAPS Expression Patterns in Mouse Cerebellum.** Heatmap displaying Spearman correlation coefficients between mouse CoGAPS-derived expression patterns (M-1 through M-13) and cerebellar cell populations including granule cells, Purkinje cells, molecular layer interneurons, interneurons, Bergmann glia, Purkinje layer interneurons, Golgi cells, and white matter. Correlation values range from high positive (red, $r = 1$) to high negative (blue, $r = -1$). Patterns demonstrate cell-type-specific enrichment, with several patterns showing preferential correlation with granule cell or Purkinje cell populations. For all analyses, $n = 2$ per group. Abbreviations: CoGAPS, Coordinated Gene Activity in Pattern Sets; GC, granule cell; Hx, hypoxia; LPS, lipopolysaccharide; M, mouse; MIA, maternal immune activation; MLI, molecular layer interneuron; N, normoxia; NES, normalized enrichment score; PC, Purkinje cell; PLI, Purkinje layer interneuron; Sal, saline; WM, white matter.

**Supplementary Fig. 4. Gene Set Enrichment Analysis of Representative CoGAPS Patterns M-1 and M-4. (A–B)** Dot plots displaying the top 25 enriched Gene Ontology and Gene Ontology Molecular Function gene sets for CoGAPS pattern M-1 (A) and pattern M-4 (B). Gene sets are drawn from Gene Ontology Cellular Component (GOCC), Gene Ontology Biological Process (GOBP), and Gene Ontology Molecular Function (GOMF) databases. Circle size represents statistical significance, expressed as $-\log_{10}(P)$, ranging from 15 to 30. Circle color represents normalized enrichment score (NES), ranging from 1.1 (light pink) to 1.5 (dark red). Pattern M-1 is enriched for gene sets associated with synaptic structure, neuron projection, somatodendritic compartment, axon, postsynapse, presynapse, synaptic signaling, cell-cell signaling, neurogenesis, neuron differentiation, dendritic tree, and vesicle-mediated transport. Pattern M-4 is enriched for gene sets associated with mitochondrial organization and structure (mitochondrion, mitochondrial envelope, mitochondrial protein-containing complex, respirasome), oxidative phosphorylation, aerobic respiration, ATP metabolic process, energy derivation by oxidation of organic compounds, generation of precursor metabolites and energy, cytosolic ribosome, and active transmembrane transporter activity. Abbreviations: CoGAPS, Coordinated Gene Activity in Pattern Sets; GOBP, Gene Ontology Biological Process; GOCC, Gene Ontology Cellular Component; GOMF, Gene Ontology Molecular Function; M, mouse; NES, normalized enrichment score.

**Supplementary Fig. 5. Insult-Specific Remodeling of Predicted Ligand-Receptor Interactions in the Cerebellar Connectome. (A–D)** LIANA plots displaying the top 20 predicted ligand-receptor interactions for each experimental group: Sal-N (A), Sal-Hx (B),

LPS-N (C), and LPS-Hx (D). For each group, interactions are organized by source cell type (granule cell or Purkinje cell, columns) and target cell type (granule cell or Purkinje cell, sub-columns), yielding four interaction categories: GC→GC, GC→PC, PC→GC, and PC→PC. Rows represent individual ligand-receptor pairs. Circle size represents interaction specificity (natmi.edge_specificity), ranging from 0.08 (small) to 0.54 (large). Circle color represents expression magnitude (sca.LRscore), ranging from 0.5 (purple, low) to 0.9 (yellow, high). For analysis, $n = 2$ per group. Abbreviations: GC, granule cell; Hx, hypoxia; LIANA, LIgand-receptor ANalysis frAmework; LPS, lipopolysaccharide; N, normoxia; PC, Purkinje cell; Sal, saline; sca.LRscore, scaled ligand-receptor score.

**Supplementary Fig. 6. Spatial Distribution of Exploratory Behavior During the Three-Chamber Social Approach Assay. (A)** Representative schematic of the six-grid spatial analysis displaying time distribution across arena regions. The arena was divided into a 2 × 3 grid (top, middle, bottom rows; left and right columns), with time spent in each grid region indicated both by color intensity ranging from 50 seconds (yellow) to 350 seconds (red) and by actual time shown as minute:seconds below clock icon. **(B)** Representative movement tracking schematics for the three-chamber social approach assay across experimental conditions. Stranger mouse location is indicated by an x-symbol; novel object location is indicated by a circle-symbol. Dashed lines represent chamber boundaries. Color scale indicates time spent per spatial bin ranging from 0.005 seconds (blue) to 0.020 seconds (red). Abbreviations: LPS, lipopolysaccharide; Sal, saline; Hx, hypoxia; N, normoxia.

**Supplementary Fig. 7. Representative Kinematic Motifs for B-SOiD-Identified Behavioral Clusters.** Representative still frames from video segments illustrating the 14 distinct kinematic clusters (Clusters 0-13) identified by the B-SOiD algorithm. Each frame exemplifies a characteristic locomotor motif corresponding to a specific data-driven behavioral cluster extracted from high-resolution movement tracking during the three-chamber social approach assay. Abbreviations: B-SOiD, Behavioral Segmentation of Open-field In DeepLabCut.

**Supplementary Fig. 8. Chamber-specific B-SOiD Kinematic Profiles Across Experimental Groups in the Three-chamber Social Approach Assay.** Heatmaps display the fraction of total frames occupied by each B-SOiD-identified kinematic cluster per experimental group. Rows correspond to individual kinematic clusters; columns represent experimental groups (Sal-N, Sal-Hx, LPS-N, LPS-Hx). Distributions are shown pooled across all chambers and decomposed by individual chamber: Stranger Mouse, Middle, and Novel Object. Color intensity encodes the fraction of frames with scale: 0 (light blue) to 0.20 (dark blue) total fraction of frames. For all analyses, *n* = 6-15 per group. Abbreviations: LPS, lipopolysaccharide; Sal, saline; Hx, hypoxia; N, normoxia.

**Supplementary Fig. 9. Chamber-Resolved Bout Duration Distributions across B-SOiD Kinematic Clusters.** Cumulative distribution function plots displaying the duration of behavioral bouts for all 14 kinematic clusters (Clusters 0-13) across the three chambers (stranger mouse chamber, middle chamber, and novel object chamber) for each experimental group. Duration is expressed as successive frames on a logarithmic scale (x-axis), with cumulative distribution function values on the y-axis. Each row represents a distinct kinematic cluster, and each column represents a chamber location. Line colors indicate experimental groups: Sal-N (green), Sal-Hx (yellow), LPS-N (blue), and LPS-Hx (red). Asterisks indicate comparisons with at least one statistically significant interaction between groups. For all analyses, *n* = 6-15 per group; Wasserstein distance with Benjamini-Hochberg correction applied to P-values. CDF, cumulative distribution function; Hx, hypoxia; LPS, lipopolysaccharide; N, normoxia; Sal, saline.

**Supplementary Table 1 Primary and secondary antibodies used for immunofluorescence**

| Antibody | Conjugate | Isotype | Dilution | Catalogue no. | Lot no. | Company | Use |
|---|---|---|---|---|---|---|---|
| VGLUT2 | Alexa Fluor 594 | Guinea pig | 1:200 (IF) | 135404 | 3-46 | Synaptic Systems | 1° |
| CB300 | Alexa Fluor 488 | Mouse monoclonal, IgG1 | 1:200 (IF) | 300 | 07(F) | SWANT | 1° |
| P27 | Alexa Fluor 594 | Rabbit polyclonal | 1:200 (IF) | Sc529 | C1313 | Santa Cruz Biotechnology, Inc. | 1° |
| P21 | Alexa Fluor 488 | Mouse monoclonal, IgG | 1:200 (IF) | Op76 | D00094288 | CalBiochem, Sigma Aldrich | 1° |
| Anti-Rabbit | Alexa Fluor 594 | Rabbit | 1:250 (IF) | 711585152 | 163459 | Jackson ImmunoResearch Laboratories, Inc. | 2° |
| Anti-Mouse | Alexa Fluor 488 | Mouse | 1:250 (IF) | 715545151 | 164101 | Jackson ImmunoResearch Laboratories, Inc. | 2° |
| Anti-Guinea pig | Alexa Fluor 594 | Guinea pig | 1:250 (IF) | 706585148 | 149524 | Jackson ImmunoResearch Laboratories, Inc. | 2° |
| Anti-Goat | Alexa Fluor 647 | Goat | 1:250 (IF) | 705605147 | 160039 | Jackson ImmunoResearch Laboratories, Inc. | 2° |

Antibodies, conjugated fluorophores, host/isotype, working dilutions, catalogue and lot numbers, and supplier for all immunostaining performed in mouse and human cerebellar tissue.

*Abbreviations: IF = immunofluorescence; 1° = primary antibody; 2° = secondary antibody.*

**Supplementary Table 2 Demographic and clinical characteristics of preterm and term human subjects included in the transcriptomics analysis**

| **Variable** | **Term (n = 4)** | | | **Preterm (n = 4)** | | |
|---|---|---|---|---|---|---|
| | **Mean ± SD** | **n** | **%** | **Mean ± SD** | **n** | **%** |
| ***Sex*** | | | | | | |
| Female | | 3 | 75 | | 3 | 75 |
| Male | | 1 | 25 | | 1 | 25 |
| ***Race*** | | | | | | |
| Black | | 2 | 50 | | 4 | 100 |
| White | | 2 | 50 | | 0 | 0 |
| ***Ethnicity*** | | | | | | |
| Hispanic or Latino | | 1 | 25 | | 0 | 0 |
| Not Hispanic or Latino | | 2 | 50 | | 3 | 75 |
| Unknown | | 1 | 25 | | 1 | 25 |
| GA[a] (weeks) | 40 ± 0 | | | 26.4 ± 0.5 | | |
| Postmenstrual age (weeks) | 46 ± 9.3 | | | 27.1 ± 1 | | |
| Brain weight (g) | 435.5 ± 82.8 | | | 126.95 ± 18.57 | | |

[a] Gestational age (GA) determined at delivery.

*Abbreviations: GA = gestational age; SD = standard deviation.*

**Supplementary Table 3 Demographic and clinical characteristics of human preterm subjects included in the transcriptomics analysis**

| Subject ID | GA[a] (weeks) | Postmenstrual age (weeks) | Sex | Brain weight (g) | Race | Ethnicity | Cause of death |
|---|---|---|---|---|---|---|---|
| Preterm 1 | 26.4 | 27 | F | 116.8 | B | NH | Prematurity, bowel perforation |
| Preterm 2 | 26 | 26.1 | F | 121.5 | B | UK | Prematurity, ELBW |
| Preterm 3 | 27.3 | 28.7 | F | 154.5 | B | NH | Prematurity, RDS, intrauterine fetal hemorrhage |
| Preterm 4 | 26 | 26.4 | M | 115 | B | NH | Bronchopneumonia |

[a] Gestational age (GA) determined at delivery.

*Abbreviations: B = Black; ELBW = extremely low birth weight; F = female; GA = gestational age; M = male; NH = non-Hispanic; RDS = respiratory distress syndrome; UK = unknown.*

**Supplementary Table 4 Demographic and clinical characteristics of human term subjects included in the transcriptomics analysis**

| Subject ID | GA[a] (weeks) | Postmenstrual age (weeks) | Sex | Brain weight (g) | Race | Ethnicity | Cause of death |
|---|---|---|---|---|---|---|---|
| Term 1 | 40 | 40.7 | F | 404.5 | W | UK | Sepsis, coagulopathy, multi-organ system failure |
| Term 2 | 40 | 41 | F | 404.3 | B | NH | HSVII |
| Term 3 | 40 | 62.1 | F | 575 | W | HL | Hypoplastic left ventricle, double outlet right ventricle |
| Term 4 | 40 | 40.3 | M | 358 | B | NH | CPT II deficiency |

[a] Gestational age (GA) determined at delivery.

*Abbreviations: B = Black; F = female; GA = gestational age; HL = Hispanic Latino; HSVII = herpes simplex virus II; M = male; NH = non-Hispanic; UK = unknown; W = White.*

**Supplementary Table 5 Demographic and clinical characteristics of the human postmortem cerebellar cohort (n = 8), stratified by gestational age**

| Characteristic | Preterm (n = 4) | Term (n = 4) |
|---|---|---|
| ***Demographics and birth*** | | |
| Gestational age, weeks, median [Q1, Q3] | 26 [26, 26.3] | 40 [40, 40] |
| Birth weight, g, median [Q1, Q3] | 845 [777.5, 890.2] | 3212.5 [2945, 3441.2] |
| Female sex, n (%) | 3 (75%) | 3 (75%) |
| Cesarean delivery, n (%) | 3 (75%) | 0 (0%) |
| Age at death, days, median [Q1, Q3] | 3.5 [2.5, 5.5] | 6 [4.8, 44] |
| ***Documented prenatal and perinatal inflammatory exposures*** | | |
| Clinical chorioamnionitis | 2 (50%) | 0 (0%) |
| Documented maternal infection during pregnancy | 2 (50%) | 0 (0%) |
| Intrapartum antibiotic administration | 1 (25%) | 0 (0%) |
| Group B Streptococcus colonization or infection | 1 (25%) | 1 (25%) |
| Elevated neonatal C-reactive protein | 3 (75%) | 1 (25%) |
| Abnormal neonatal absolute neutrophil count or I:T ratio | 3 (75%) | 2 (50%) |
| ***Documented perinatal and postnatal hypoxic exposures*** | | |
| Low Apgar score (1- or 5-minute) | 2 (50%) | 0 (0%) |
| Delivery-room resuscitation beyond initial steps | 4 (100%) | 0 (0%) |
| Mechanical ventilation initiated within 72h of birth | 4 (100%) | 2 (50%) |
| Supplemental oxygen requirement on postnatal days 3–7 | 3 (75%) | 4 (100%) |
| Sustained or recurrent desaturation events in first 72h | 2 (50%) | 2 (50%) |

Continuous variables are reported as median [Q1, Q3]; categorical variables as n (%). Cases were stratified by gestational age (preterm: <37 weeks; term: ≥37 weeks). Clinical exposures were documented from medical records and are presented to characterize the inflammatory and hypoxic exposures of the cohort.

**Supplementary Table 6 Sampling for electron-microscopy mitochondrial morphometric analysis**

| Cell type | Group | Age | No. of samples | No. of cells | No. of mitochondria |
|---|---|---|---|---|---|
| Granule | Sal-N | P11 | 4 | 15 | 57 |
| Granule | Sal-Hx | P11 | 3 | 15 | 90 |
| Granule | LPS-N | P11 | 3 | 17 | 114 |
| Granule | LPS-Hx | P11 | 4 | 20 | 102 |
| Purkinje | Sal-N | P11 | 4 | 6 | 757 |
| Purkinje | Sal-Hx | P11 | 3 | 4 | 306 |
| Purkinje | LPS-N | P11 | 4 | 6 | 712 |
| Purkinje | LPS-Hx | P11 | 4 | 8 | 694 |
| Granule | Sal-N | P45 | 3 | 19 | 77 |
| Granule | Sal-Hx | P45 | 2 | 10 | 42 |
| Granule | LPS-N | P45 | 2 | 10 | 41 |
| Granule | LPS-Hx | P45 | 4 | 20 | 57 |
| Purkinje | Sal-N | P45 | 4 | 10 | 1783 |
| Purkinje | Sal-Hx | P45 | 2 | 4 | 682 |
| Purkinje | LPS-N | P45 | 2 | 3 | 528 |
| Purkinje | LPS-Hx | P45 | 3 | 5 | 863 |

Number of biological samples, segmented cells, and individual mitochondria analyzed per cell type, experimental group, and age in the two-dimensional mitochondrial segmentation analysis.

*Abbreviations: Sal = saline; LPS = lipopolysaccharide; N = normoxia; Hx = hypoxia; P = postnatal day.*

**Supplementary Table 7 Group-specific sample sizes for each behavioral assay**

| Group | Locomotor assay (Fig. 1C–D) | SIT (Fig. 7A–B) | OFT (Fig. 7C–D) | B-SOiD (Fig. 7E–F) |
|---|---|---|---|---|
| Sal-N | 10 | 15 | 5 | 15 |
| Sal-Hx | 6 | 6 | 6 | 6 |
| LPS-N | 6 | 13 | 5 | 13 |
| LPS-Hx | 6 | 10 | 8 | 10 |
| Total | 28 | 44 | 24 | 44 |

Number of mice analyzed per experimental group in each behavioral assay after assay-specific quality control. Corresponding main-text figure panels are indicated in the column headers.

*Abbreviations: SIT = Social Interaction Test; OFT = Open Field Test; B-SOiD = Behavioral Segmentation of Open-field In DeepLabCut; Sal = saline; LPS = lipopolysaccharide; N = normoxia; Hx = hypoxia.*

**Supplementary Table 8 B-SOiD kinematic motif classification**

| Cluster | Behavioral label | Detailed description |
|---|---|---|
| 0 | Standing | Stationary posture with minimal body or head movement, with no overt locomotor or investigatory activity. |
| 1 | Left head turn with body reorientation | Leftward head turn accompanied by reorientation of the body axis, with minimal forward displacement. |
| 2 | Approaching and turning | Forward movement toward a target or location followed by body reorientation. |
| 3 | Standing and exploring | Stationary body posture accompanied by active head movements and local sensory investigation. |
| 4 | Standing with head turn | Stationary posture with discrete head rotations while the body remains largely fixed. |
| 5 | Grooming | Self-directed maintenance behavior, including licking, scratching, or cleaning of body parts. |
| 6 | Sniffing | Active olfactory investigation of the environment, substrate, object, or conspecific, characterized by repeated nose-directed movements. |
| 7 | Approaching forward | Sustained forward locomotion toward a location, object, or stimulus, with limited lateral reorientation. |
| 8 | Standing, approaching, and exploring | Mixed behavioral state combining brief stationary observation, short forward advances, and local sensory investigation. |
| 9 | Head turn | Isolated head reorientation while the body remains stationary, without accompanying locomotor displacement. |
| 10 | Turning around / circle back | Large-scale body reorientation involving reversal of movement direction or a partial circular trajectory. |
| 11 | Right body turn | Rightward reorientation of the body axis with minimal forward displacement. |
| 12 | Right body turn with forward movement | Rightward body rotation combined with forward locomotion. |
| 13 | Disengaging | Movement away from a standing position on all four limbs, accompanied by backward turning away from the current investigatory posture. |

*Abbreviations: B-SOiD = Behavioral Segmentation of Open-field In DeepLabCut.*

**Supplementary Table 9 ErasmusLadder two-way repeated-measures ANOVA with Tukey HSD post hoc comparisons (Session 5)**

**A. Post-perturbation step-time**

| Two-way RM ANOVA[a] | SS | DF | MS | F (DFn, DFd) | P value |
|---|---|---|---|---|---|
| Sessions × Condition | 542830 | 21 | 25849 | F (21, 168) = 1.808 | P = 0.0212 |
| Sessions | 2297170 | 7 | 328167 | F (7, 168) = 22.96 | P < 0.0001 |
| Condition | 93946 | 3 | 31315 | F (3, 24) = 1.081 | P = 0.3760 |
| Subject | 695348 | 24 | 28973 | F (24, 168) = 2.027 | P = 0.0052 |
| Residual | 2401459 | 168 | 14294 | | |

**Tukey's multiple comparisons test — Session 5**

| # | Tukey's multiple comparisons test[b] | Predicted (LS) mean diff. | 95% CIs | Summary | Adjusted P value |
|---|---|---|---|---|---|
| 5 | Sal-N Post-perturbation vs. Sal-Hx Post-perturbation | −186.57 | −316.00 to −57.14 | * | 0.0254 |
| 5 | Sal-N Post-perturbation vs. LPS-N Post-perturbation | −219.75 | −349.18 to −90.32 | ** | 0.0054 |
| 5 | Sal-N Post-perturbation vs. LPS-Hx Post-perturbation | −366.07 | −495.50 to −236.64 | **** | < 0.0001 |
| 5 | Sal-Hx Post-perturbation vs. LPS-N Post-perturbation | −33.18 | −177.89 to 111.52 | ns | 0.9691 |
| 5 | Sal-Hx Post-perturbation vs. LPS-Hx Post-perturbation | −179.50 | −324.21 to −34.79 | ns | 0.0720 |
| 5 | LPS-N Post-perturbation vs. LPS-Hx Post-perturbation | −146.32 | −291.02 to −1.61 | ns | 0.1936 |

**B. Missteps**

| Two-way RM ANOVA | SS | DF | MS | F (DFn, DFd) | P value |
|---|---|---|---|---|---|
| Sessions × Condition | 623.3 | 21 | 29.68 | F (21, 168) = 1.694 | P = 0.0359 |
| Sessions | 878.3 | 7 | 125.5 | F (7, 168) = 7.161 | P < 0.0001 |
| Condition | 283.5 | 3 | 94.49 | F (3, 24) = 0.9169 | P = 0.4476 |
| Subject | 2473 | 24 | 103.1 | F (24, 168) = 5.881 | P < 0.0001 |
| Residual | 2944 | 168 | 17.52 | | |

**Tukey's multiple comparisons test — Session 5**

| # | Tukey's multiple comparisons test | Predicted (LS) mean diff. | 95% CIs | Summary | Adjusted P value |
|---|---|---|---|---|---|
| 5 | Sal-N vs. Sal-Hx | −10.03 | −15.47 to −4.58 | ** | 0.0023 |
| 5 | Sal-N vs. LPS-N | 1.89 | −3.55 to 7.34 | ns | 0.9006 |
| 5 | Sal-N vs. LPS-Hx | 1.19 | −4.25 to 6.64 | ns | 0.9724 |
| 5 | Sal-Hx vs. LPS-N | 11.92 | 5.83 to 18.01 | ** | 0.0011 |
| 5 | Sal-Hx vs. LPS-Hx | 11.22 | 5.13 to 17.30 | ** | 0.0023 |
| 5 | LPS-N vs. LPS-Hx | −0.70 | −6.79 to 5.39 | ns | 0.9958 |

[a] Two-way repeated-measures ANOVA performed on Sessions 1–8 with Condition (Sal-N, Sal-Hx, LPS-N, LPS-Hx) as the between-subjects factor and Session as the within-subjects factor; n = 7 per group.

[b] Tukey's HSD post hoc multiple-comparisons test applied at Session 5 (perturbation session). 95% CIs are presented for the predicted (least-squares) mean differences.

*P < 0.05, **P < 0.01, ***P < 0.001, ****P < 0.0001; ns, not significant. Bold marks statistically significant comparisons.

NOTE: Please save this file locally before filling in the table, DO NOT work on the file within your internet browser as changes will not be saved. Adobe Acrobat Reader (available free here) is recommended for completion.

### The ARRIVE guidelines 2.0: author checklist

#### The ARRIVE Essential 10

These items are the basic minimum to include in a manuscript. Without this information, readers and reviewers cannot assess the reliability of the findings.

| Item | | Recommendation | Section/line number, or reason for not reporting |
|---|---|---|---|
| **Study design** | 1 | For each experiment, provide brief details of study design including: | |
| | | a. The groups being compared, including control groups. If no control group has been used, the rationale should be stated. | Methods, mouse model. Four groups: Sal-N (control), Sal-Hx, LPS-N, LPS-Hx. |
| | | b. The experimental unit (e.g. a single animal, litter, or cage of animals). | Methods, mouse model. Experimental unit = individual animal. |
| **Sample size** | 2 | a. Specify the exact number of experimental units allocated to each group, and the total number in each experiment. Also indicate the total number of animals used. | Methods + figure legends. n per group/assay reported (histology 4-5; Seahorse 5-14; behaviour 24-44; spatial 2; human 4+4). |
| | | b. Explain how the sample size was decided. Provide details of any *a priori* sample size calculation, if done. | Methods, statistics. Resource-equation approach; no a priori power calculation. |
| **Inclusion and exclusion criteria** | 3 | a. Describe any criteria used for including and excluding animals (or experimental units) during the experiment, and data points during the analysis. Specify if these criteria were established *a priori*. If no criteria were set, state this explicitly. | Methods, behaviour + cross-species. Pre-set criteria: tracking/recording quality, >25% jumps, ROUT (Q=1%), PCA outliers. |
| | | b. For each experimental group, report any animals, experimental units or data points not included in the analysis and explain why. If there were no exclusions, state so. | Methods + legends. |
| | | c. For each analysis, report the exact value of *n* in each experimental group. | Exact n per group in Methods tables and figure legends. |
| **Randomisation** | 4 | a. State whether randomisation was used to allocate experimental units to control and treatment groups. If done, provide the method used to generate the randomisation sequence. | Methods, mouse model. Randomised at two levels via R; sequence by investigator blind to outcomes. |
| | | b. Describe the strategy used to minimise potential confounders such as the order of treatments and measurements, or animal/cage location. If confounders were not controlled, state this explicitly. | Methods, mouse model. Litters distributed across groups; litter treated as biological variability. |
| **Blinding** | 5 | Describe who was aware of the group allocation at the different stages of the experiment (during the allocation, the conduct of the experiment, the outcome assessment, and the data analysis). | Methods, blinding. Coded/decoded after quantification; all scoring blind; hypoxia exposure could not be blinded. |
| **Outcome measures** | 6 | a. Clearly define all outcome measures assessed (e.g. cell death, molecular markers, or behavioural changes). | Methods + Results. Motor (missteps, step-time), histology, VGLUT2, p21/p27, OCR, mitochondria, cell cycle, social/open-field, B-SOiD motifs. |
| | | b. For hypothesis-testing studies, specify the primary outcome measure, i.e. the outcome measure that was used to determine the sample size. | Methods, statistics. Exploratory; no single pre-specified primary outcome. |
| **Statistical methods** | 7 | a. Provide details of the statistical methods used for each analysis, including software used. | Methods, statistics. Prism/R. ANOVA (Tukey/Fisher LSD), Kruskal-Wallis/Dunn, RM-ANOVA, Mann-Whitney U, Welch t, Wilcoxon, Wasserstein/BH. |
| | | b. Describe any methods used to assess whether the data met the assumptions of the statistical approach, and what was done if the assumptions were not met. | Methods, statistics. Shapiro-Wilk (normality) + Levene (variance); non-parametric fallback when violated. |
| **Experimental animals** | 8 | a. Provide species-appropriate details of the animals used, including species, strain and substrain, sex, age or developmental stage, and, if relevant, weight. | Methods, mouse model. C57BL/6 (#000664) and CD1; both sexes (counts given); P3-P45. |
| | | b. Provide further relevant information on the provenance of animals, health/immune status, genetic modification status, genotype, and any previous procedures. | Methods, mouse model. Wildtype; standard housing; hypoxic pups cross-fostered to CD1; no prior procedures. |
| **Experimental procedures** | 9 | For each experimental group, including controls, describe the procedures in enough detail to allow others to replicate them, including: | |
| | | a. What was done, how it was done and what was used. | Methods. LPS IP (33 ug/mL, 4.5 uL/g) E15-16; hypoxia 10.5-11% FiO2; histology, EM, Seahorse, Visium, behaviour. |
| | | b. When and how often. | Methods. MIA E15-16; hypoxia P3-P11; Erasmus P35-42; social P44; tissue P11 and P45. |
| | | c. Where (including detail of any acclimatisation periods). | Methods, behaviour. At Children's National Hospital; light phase; 20-30 min acclimatisation; foster dams in chamber. |
| | | d. Why (provide rationale for procedures). | Methods. LPS E15-16 = 22-24 wk human; hypoxia from P3 (P1-2 lethal); validated paradigm; CD1 fostering rationale given. |
| **Results** | 10 | For each experiment conducted, including independent replications, report: | |
| | | a. Summary/descriptive statistics for each experimental group, with a measure of variability where applicable (e.g. mean and SD, or median and range). | Results + legends. Group summary stats with variability reported. [On box-plot conversion, report median + IQR.] |
| | | b. If applicable, the effect size with a confidence interval. | Methods + Suppl. Fig. 1. AUC with 95% CI for cross-species enrichment; CIs not computed elsewhere. |
